# Investigation and optimization the effect of electrical stimulation parameters on the differentiation of human adipose mesenchymal stem cells into neurons-like cells on carbon nanofibers

**DOI:** 10.1101/2024.05.08.593090

**Authors:** Houra Nekounam, Hossein Golmohammadi, Seyed Mohammad Amini, Mohammad Ali Shokrgozar, Reza Faridi-Majid

**Affiliations:** Department of Medical Nanotechnology, School of Advanced Technologies in Medicine, Tehran University of Medical Sciences, Tehran, Iran; Department of Engineering, Islamic Azad University of South-Tehran Branch, Tehran, Iran; Radiation Biology Research Center, Iran University of Medical Sciences, Tehran, Iran; National Cell Bank of Iran, Pasteur Institute of Iran, Tehran, Iran

**Keywords:** Neural tissue engineering, Carbon nanofibers, Electro-conductive Scaffolds, Electrical Stimulation

## Abstract

**Background:** Neurodegenerative diseases are among the most challenging diseases because neuron cells are not able to regenerate spontaneously. Tissue engineering is one of the most promising stem cell-based therapies. Controlling stem cell differentiation is a very crucial aspect of tissue engineering.

**Methods:** In this study, carbon nanofibers with an average diameter of 181±45 nm were prepared as a conductive scaffold based on the electrospinning method and subsequent thermal processing. Scaffold structure characterization were performed with XRD, Raman and Electrical conductivity tests. A homemade device was prepared to transmit electrical current to cells seeded on the scaffold in a culture plate. Various current parameters such as current intensity, frequency, waveform, daily shock duration, and shock period on adipose mesenchymal stem cells were examined for differentiation into neuronal cells. SPSS software and the one-way analysis of variance (ANOVA) was used as statistical analysis.

**Results:** Characterization tests confirmed the formation of the carbon and crystallite structure with the electrical conductivity . Current with 1500 uA intensity, 500Hz frequency, and square waveform were selected as the optimal current parameters. It was found that the daily and periodic increase in shock time leads to an increase in the expression of neural and glial genes. A comparison of groups with real-time PCR and immunofluorescence of nestin, Map2, TubB3, and GFAPgenes was evaluated.

**Conclusions:** There are a variety of chemical and physical methods to control cell behavior, one of which is electrical stimulation. Conductive scaffolding is required for direct electrical stimulation of cells. The results showed that the method based on electrical stimulation can well cause neural differentiation, and considering the problems in preparing and maintaining chemical differentiation agents, it can be used practically.

**Graphical abstract:**
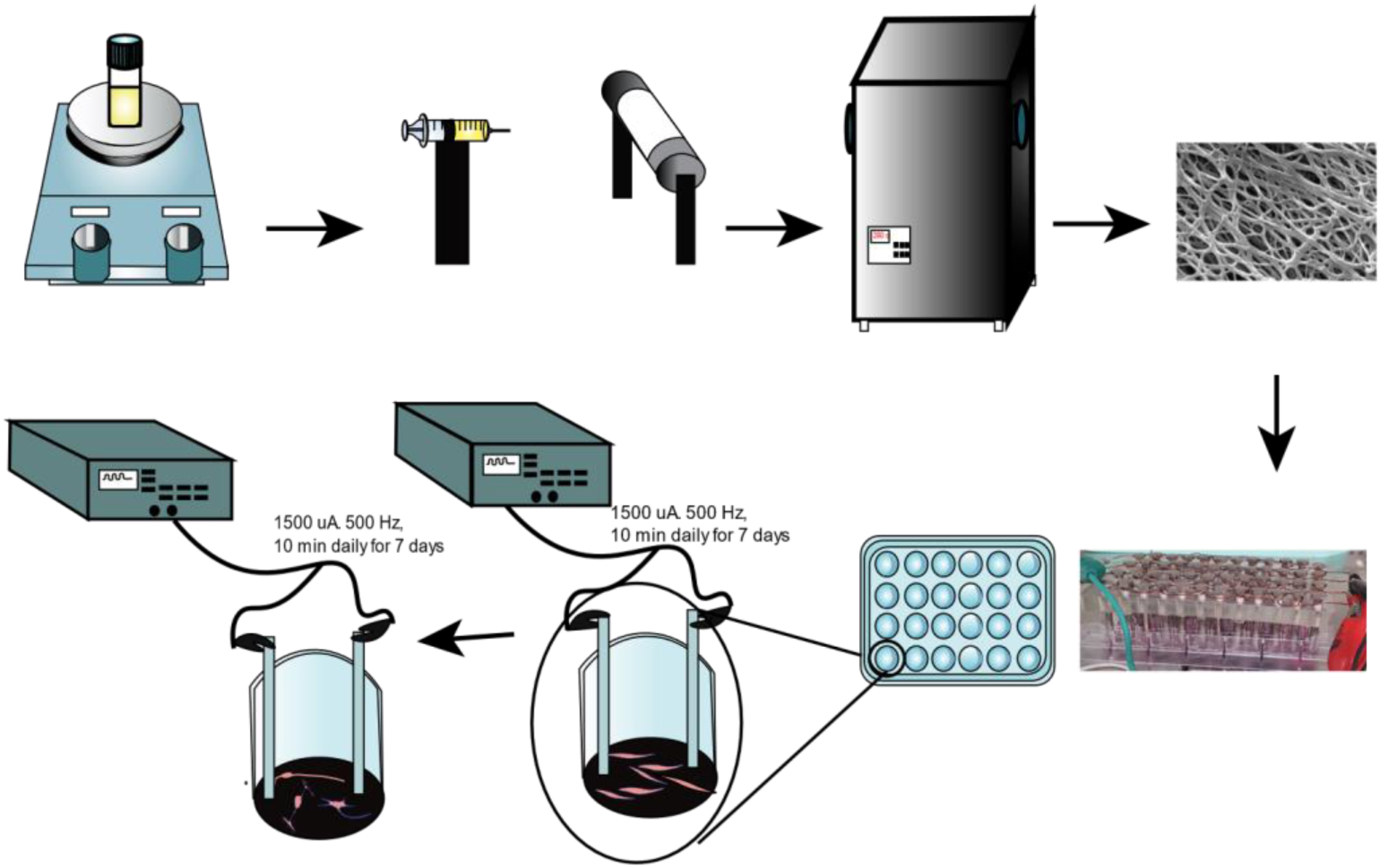
stages of scaffold preparation and electrical stimulation of stem cells

## Background

The increase in population and the increase in the average age of the world population will lead to a definite increase in neurodegenerative diseases, injuries caused by accidents, or other injuries of the nervous system. Even nowadays, degenerative nerve diseases, including Alzheimer’s, Parkinson’s, and stroke affect a high percentage of the population[1–3]. The conventional treatment of these diseases due to the complexity of the nervous system is challenging. The treatment of nerve degenerative diseases is difficult, time-consuming, expensive, and inefficient for functional recovery and nerve regeneration [4]. Therefore, providing new solutions to improve the function of damaged parts of the nervous system is one of the most important research in tissue engineering [5]. The differentiation of stem cells into nerves and glial cells can be a useful approach for recovery function, secreting cytokines, and growth factors, and inhibiting cell apoptosis or fibrosis via endogenous repair activated by immunomodulation [4]. Presenting techniques for the differentiation of stem cells in prosthetic scaffolds is a crucial component of nervous system tissue engineering. These emerging and interdisciplinary methods provide improved and new scaffolding with cells that can improve, maintain, and regenerate neural tissue function [3].

Current chemical techniques for neural differentiation, suffer from many limitations due to chemical reagents and small molecules’ unwanted effects. For example, some substances currently applied for neuronal differentiation, such as dimethylsulfoxide (DMSO), b-mercaptoethanol (BME), and butylated Hydroxyanisole (BHA) are restricted due to cytotoxicity and induced cellular stress [5, 6]. So cell differentiation using non-chemical methods has attracted much attention [4].

Various biochemical and biophysical signs guide the development, and maintenance of the structure and function of live organisms. Therefore, understanding and using these stimuli to improve the quality of tissue engineering implants is desirable. Chemical, mechanical, material-based (i.e., topography, scaffolds), and magnetic cues have been used to increase efficiency and improve in vitro tissue preparation parameters[7–9].

The regulation of the physical and chemical properties of biomaterials in tissue engineering is crucial to creating functional properties. Electrical stimulation combined with electrical conductive biomaterials that are biocompatible increases the biological effect of electrical stimulation [10]. The primary source of body electricity is the cells due to the continuous pumping of ion channels, which creates the potential gradient across the membrane[11]. The cells next to each other form a resistant layer parallel to the cell membrane at a larger scale [12–14]. The endogenous electric field plays an important role in the function of all living organisms and physiological processes[7–9], including cell migration, proliferation, differentiation, and wound healing[5, 15, 16]. The electric field causes the reconstruction and repair of various tissues, including the nerve, bone, and epidermis[17–19]

Electrical stimulation is flexible, non-chemical, and can be used for in vitro and in vivo research [20]. The frequency, duration time, voltage, and current of electrical stimulation vary depending on the cell type and purpose of the study. This method can regulate the body’s potential along with physiological functions, including molecular transmission, signal transmission, and embryonic development through ion channels. Electrical stimulation also triggers physiological activities such as proliferation, differentiation, and migration of different cells [21–23]. Exogenous electrical stimulation increases migration, differentiation, neural growth, and intracellular Ca^2+^ dynamics in the extracellular environment of neuronal stem cells [24–27].

Biophysical changes begin at the cell surface, altering the function of membrane proteins including enzymatic activity, protein receptor complexes, and ion transport channels by altering the ion distribution [28]. The cellular effect of Electrical stimulation, including microfilament rearrangement that causes rearrangement of charged cell surface receptors, alters cell shape, and releases signaling molecules, and interacellular calcium. Finally, electrical stimulation affects cell proliferation, differentiation, and apoptosis[29–33]. The electric field has an important effect on directional cell migration and differentiation, including microfilament recombination, cell surface receptor rearrangement (CSR), and changes in intracellular Ca^2+^ dynamics[28, 31, 34].

Electroactive biomaterials include conductive polymers (CPs), metal nanoparticles, and carbon-based materials[35]. Annealed carbon nanofiber scaffolds combined with biphasic electrical stimulation enhance NSC proliferation, increase the expression level of neuronal genes, and increase MAP2 immunofluorescence, which improves nerve differentiation and maturation [35, 36] Nanofiber structures are useful in nerve tissue engineering because of their high porosity, high surface-to-volume ratio, and high spatial interconnectivity. the morphology of nanofibers is similar to fibrous proteins of extracellular matrix [37, 38]. Recently, carbon-based structures have been considered in neural tissues because of their electrical conductivity, excellent mechanical properties, structural similarity to neural tissue, and biocompatibility [1, 2, 39]. In one study it was observed that neuroblastoma cells and Schwann cells were compatible with carbon-derived nanofibers. Compared to flat carbon films, carbon nanofiber films show more adhesion and cell proliferation. And is said to be used as a nerve implant [40].

Many different types of stem cells have been identified, including NSCs, mesenchymal stem cells (MSCs), induced pluripotent stem cells (iPSCs), and embryonic stem cells (ESCs) have the potential for stem cells [41–45]. Adipose tissue is one of the most available sources of stem

cells that have a high proliferative capacity, multilineage potential, and Pluripotency. adipose stem cells can differentiate into nerve tissue [46–51]

## Materials and Methods

### 1.1. Materials

Polyacrylonitrile (PAN, MW=150,000 gmol^−1^, Polyacryl, Iran), Dimethylformamide (DMF-Merck), DMEM/High Glucose culture media, Fetal Bovine Serum (FBS) was purchased from Gibco (USA), Cytotoxicity Detection Kit PLUS, Roche company, Germany, (LDH) was used. MTT (3-(4,5-dimethylthiazol-2-yl)-2,5-diphenyltetrazolium bromide) and collagenase type 1 was purchased from Sigma-Aldrich. RNA extraction kit (Iran), cDNA Synthesis kit, Real-time Mastermix kit from (Biosystem, England),

### 1.2. Preparation and characterization of carbon nanofibers

Carbon nanofiber is prepared according to the protocol mentioned in our earlier articles [52, 53]. The Microstructure, morphology, and diameter of the fibers were seen by scanning electron microscopy Philips-XL30 at a voltage of 26 kV. Image j v1.50e software was used to determine the diameter of nanofibers and the distribution diagram of nanofiber diameter. Raman spectroscopy is used to observe the crystallinity, and disorder in carbon microstructures. the TEKSAN Raman spectrometer (TAKram P50C0R10) was used with a laser wavelength of 532 nm and a spectral resolution of 6 cm X-ray diffraction analysis to investigate the crystallinity of CNFs structure. The dispersion pattern of CNFs was obtained using a PHILIPS diffractometer, model PW1730, with Cu-Kα radiation (λ=1.54 °A). The electrical resistance and conductivity of thin layers are measured with a four-point probe(Signatone SYS-301). To evaluate hydrophilicity, the angle between the scaffold surface and the water drop based on LAmbda is evaluated. The angle and hydrophilicity were measured using a Mehrtav device(Iran).

### 1.3. Biological studies

#### Cell Viability evaluation by indirect MTT assay

Since carbon materials can absorb the color of formazan, it leads to a false-positive result. Therefore, the indirect method based on the ISO 10993/12 protocol was used. 7000 the hAMSC cells seed on the 24-well plate. A scaffold extract medium was added to the cells, and after 24 hours MTT cytotoxicity assessment was performed. Finally, by obtaining optical density(OD) at 570 and 630 using an ELISA reader (Elx 808 Biotech, USA), cell survival can be achieved. According to equation 1, the cell viability is compared to the control.

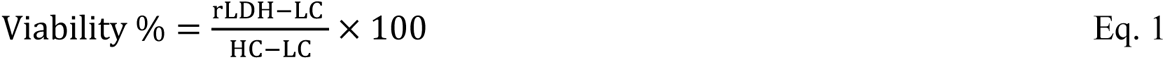

#### Direct Cytotoxicity and proliferation assay by LDH Cytotoxicity kit

CNFs were placed in a 48-well plate, and sterilization was performed through 70% ethanol (30 min) and ultraviolet light (45 min). To evaluate the cytotoxicity in the time intervals of 24, 48, and 72. The number 7000, 5000, and 3000 cells cultured per well, respectively. The cytotoxicity percent was assessed by Cytotoxicity Detection Kit plus (LDH) Roche according to Eq 2.

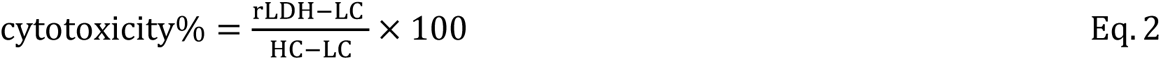

Low control contains the supernatant of cells that were cultured on the tissue culture plastic (TCP), and the high control contains the lysed cells that were cultured on the TCP. For proliferation assessment (Eq3), the cells cultured as control and on CNFs are lysed and then the amount of lactate released is measured and placed in equation 3.

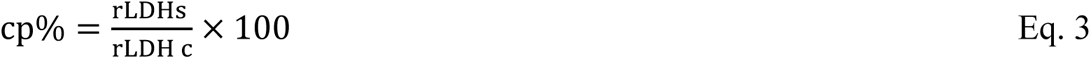

#### Isolation, culture, and Characterization of hAMSC

Stem cells were isolated according to the enzymatic extraction protocol of the adipose tissue sample from the abdominoplasty surgery of young patients aged 20-35 (Research Ethics Committees Certificate: IR.TUMS.MEDICINE.1400.1531). Flow cytometry used for immunophenotype confirmation of mesenchymal stem cells. antibodies of CD90, CD105 used as positive marker, CD34, CD45(BioLegend) used as negative marker. In addition, hAMSC cells at the third or fourth passage were cultured with osteogenic differentiation media. osteogenic media contain DEMEM/F12, 10% FBS, 1% penicillin/streptomycin, dexamethasone (10 ^-7^M), b-glycerophosphate (10mM), and ascorbic acid (50 ug/ml). The media was changed every two days. After three weeks, the fixed cells were stained with alizarin red. Another model examines the ability of cells to differentiate into fat. So 10^5^ hAMSC are cultured with adipogenic differentiation media containing DEMEM/F12; 10% FBS; 1% penicillin/streptomycin, dexamethasone (250nM); insulin (66nM); 3-isobutyl-1-methylxanthine (0.5 mM); and indomethacin (0.2 mM) add to plate. After three weeks oil-red staining was used to evaluate the adipocyte differentiation of fat-derived mesenchymal stem cells.

#### Transferring electrical current to the cells

The homemade device is designed to transmit electrical current from the generator signal to the cultured cells on the plate or scaffold. The electrodes that were in contact with the cells were selected from 316 steel, which is not toxic to the cells, and their electrical conductivity. The distance between the holes as the position of input and output electrodes in each plate well is the same. Thick plexiglass has been used as a holder for the electrodes. The plate holding the electrodes can be autoclaved and is insulated from electric current. The output and input electrodes were connected with copper wires so that the cells were in the same condition. The structure can be used for 24, 12, and 6-well plates. CNF scaffolds were placed at the bottom of the wells. The cells were cultured on the scaffold. The designed structure is easy to use and can be autoclaved daily.

### 1.4. Primer design

One of the methods to confirm neural differentiation in stem cells investigate by expression of neural genes. For this purpose, we selected famous neuron cell markers including GFAP, Tub B3, Map2, and nestin, and designed direct and reverse primer sequences for these genes. Table 1 shows the sequence of designed primers. We obtained the gene sequence from the NCBI website. Forward and reverse primers were designed using Runner Gene, 7Oligo software. GAPDH was selected as the housekeeping gene.

**Table 1:**
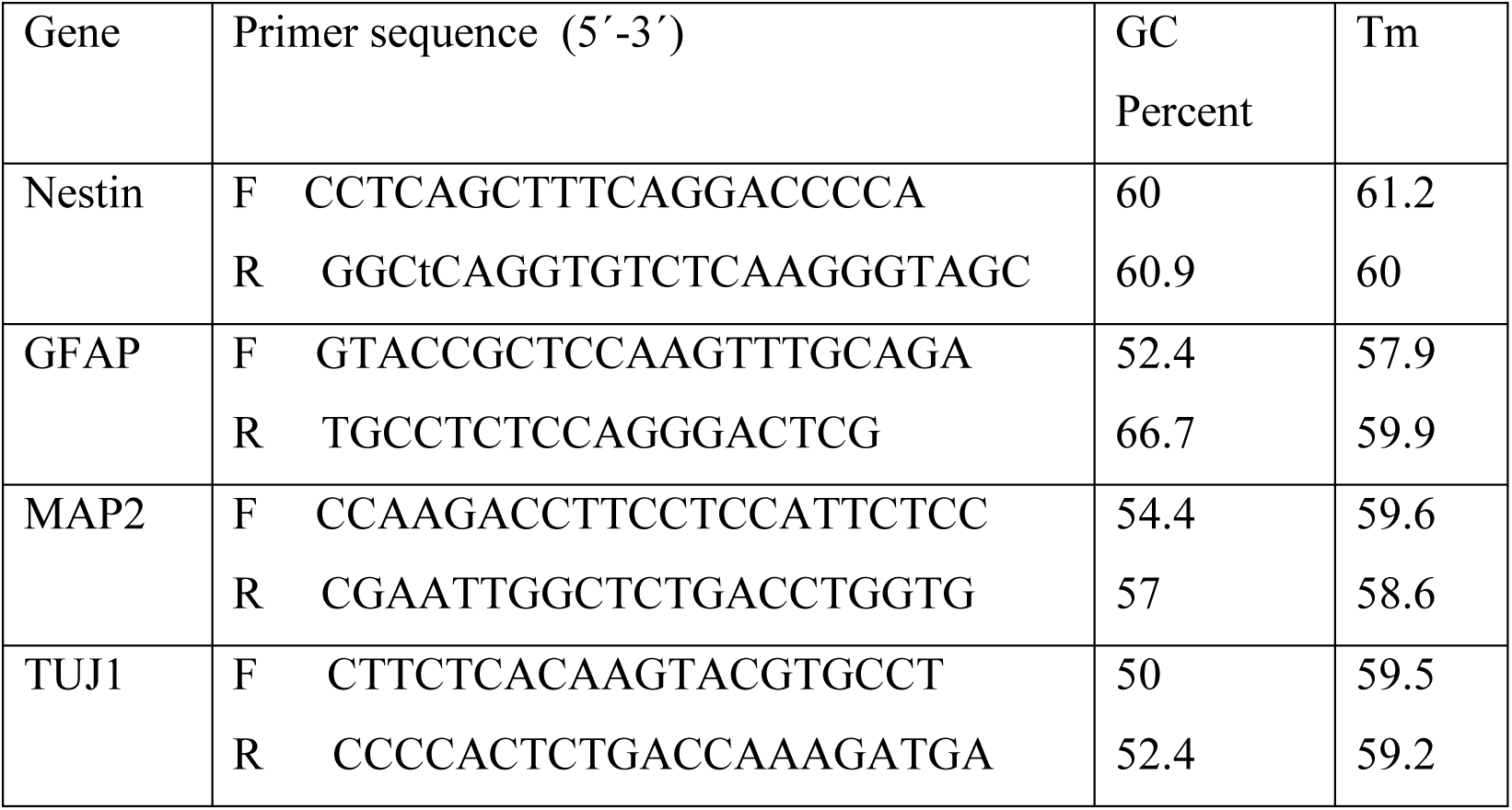
Sequences of neuronal gene primers.

### 1.5. Electrical stimulation of hAMSCs through CNFs

In the initial model, the current intensity was variable, and the rest of the factors (frequency, Waveform, daily time, and duration) were constant. The frequency waveform of the current was 100 Hz, and its waveform was the CMOS. This electric current model was repeated for seven days in 10 minutes. We tested 50, 100, 300, 600, 800, 1500, and 2000 uA, selected the appropriate current at this stage, then the frequencies of, 100, 500, and 1000 Hz were tested on the optimal current intensity. Waveform parameters were also investigated. The sinusoidal, CMOS, and square waveforms on the optimal current intensity and frequency of the previous steps were investigated. A daily time of electrical stimulation containing 10, 30, and 60 min was checked. The duration of stimulation was evaluated at 7, 14, and 21 days. Cell cultured on TCP by electrical stimulation selected as control. In the other model, the cells were evaluated simultaneously with the CNF with a neural differentiating medium. The results were evaluated by immunofluorescence and real-time PCR. For immunofluorescence, nestin, Tub B3, and Map2 markers were examined and Real-time PCR examined four neuronal markers nestin, Tub B3, MAP2, and GFAP.

### 1.6. Differentiation of stem cells by neural differentiation medium (Chemical)

The stem cells were in the passage 3-5 and healthy in terms of proliferation and appearance. The cells were cultured on scaffolds and wells (without scaffolding). The wells were filled with DMEM + 10% FBS medium to allow the cells to grow 80%. The media was is replaced with Pre-differentiation medium(.DEMEM / F12-20% FBS , 2% B27, 10 ng / ml fibroblast growth factor2, 250 μM isobutylmethylxanthin, 100 μM 2-metcaptoethanol) for 24 hours. Then neural differentiation media( DEMEM / F12, 0.2% B27, 1 μM retinoic acid (RA)/DMSO) was added to the cells for 7 days. The media was changed every two days. After 7 days, the cells entered the third medium. The third medium(DEMEM / F12, 200 ng/ml brain-derived neurotrophic factor (BDNF)) was added for another 7 days to increase cell viability. Negative control of plate cultured cells with DEMEM /10% FBS was used for the same period. The expression of differentiating factors related to neurons was examined by immunofluorescence at the level of protein expression and by real-time PCR at the level of mRNA expression.

### 1.7. Statistical analysis

The experiments were done in triplicate and the obtained data were reported as mean ± standard deviation (SD). SPSS software and the one-way analysis of variance (ANOVA) was used as statistical analysis.

## 2. Results

### 2.1. CNF preparation and characterizations

Carbon nanofiber scaffolds were electrospuned with 9% PAN/DMF solution and subjected to two thermal processes. The morphology of nanofibers was observed by SEM. In the electrospinning stage, the fibers are completely independent. During the thermal process, cross-linking occurs between the fibers and it is seen as a network of fibers. The size of the diameter of the fibers in the electrospinning stage is 120±28 nm, which can be seen in the thermal stage of the increase in the diameter of the nanofibers and reached the diameter of 181±45 nm. Few beads are recognized in the provided micrographs (Figure 1).

**Figure 1:**
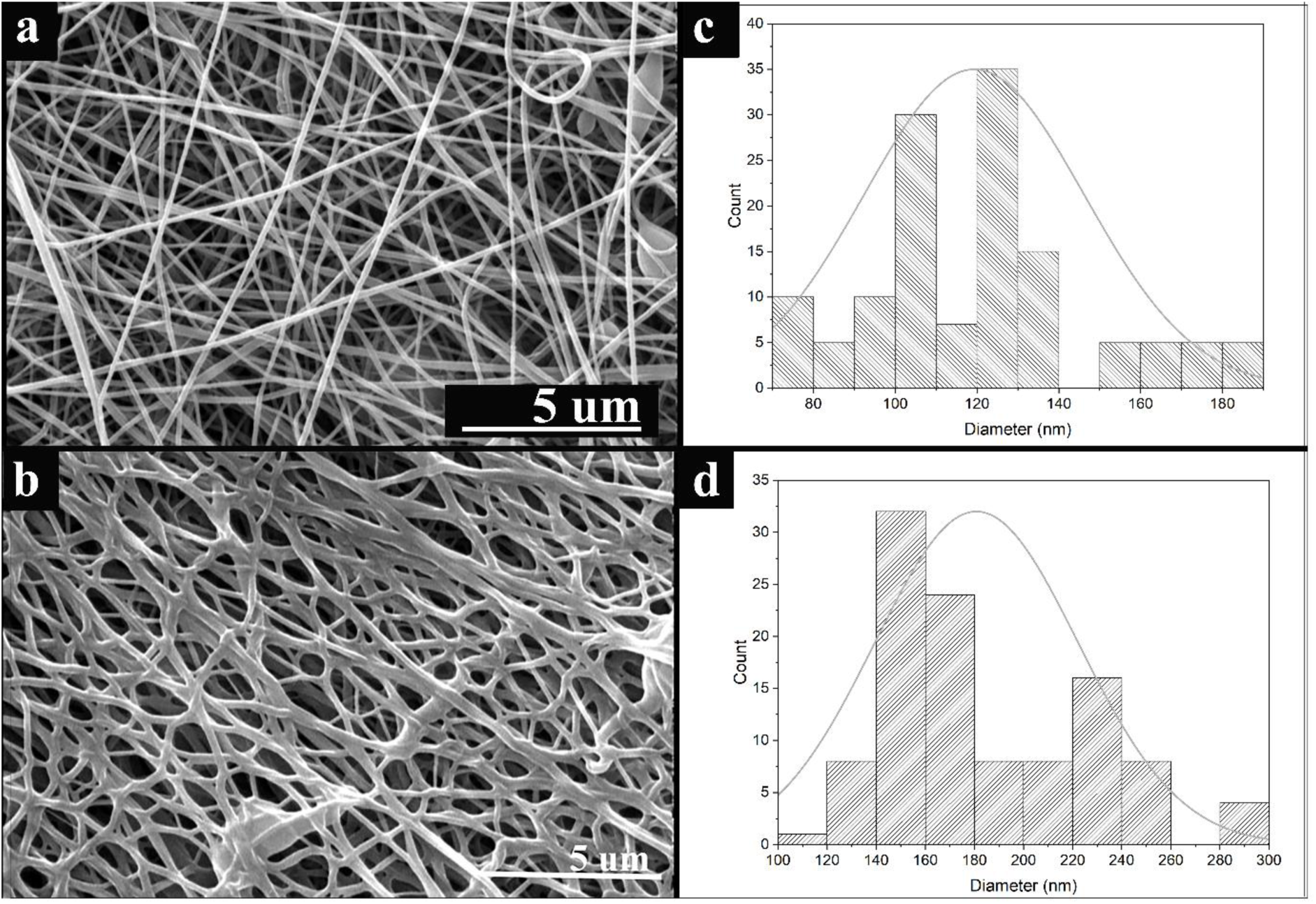
SEM and size distribution of nanofibers a, b) nanofibers electrospun and size distribution and c, d) carbon nanofibers and size distribution.

XRD results show the structure crystal of carbon nanofiber, which is characterized by the presence of the peak in regions 26 and 44. These peaks were demonstrated in Fig 2a, and the numerical value of the structures formed was calculated based on the formula that has been presented in Fig. 2(a/1). The state of structural order and disorder of the resulting graphite structure in CNF was investigated by Raman. The presence of peaks D, and G, (Fig. 2b) confirms the carbon-based structures as mentioned in our previous articles [52–56]. The calculations were performed to measure the order and disordered part, as demonstrated in Fig 2(b/1).

**Figure 2:**
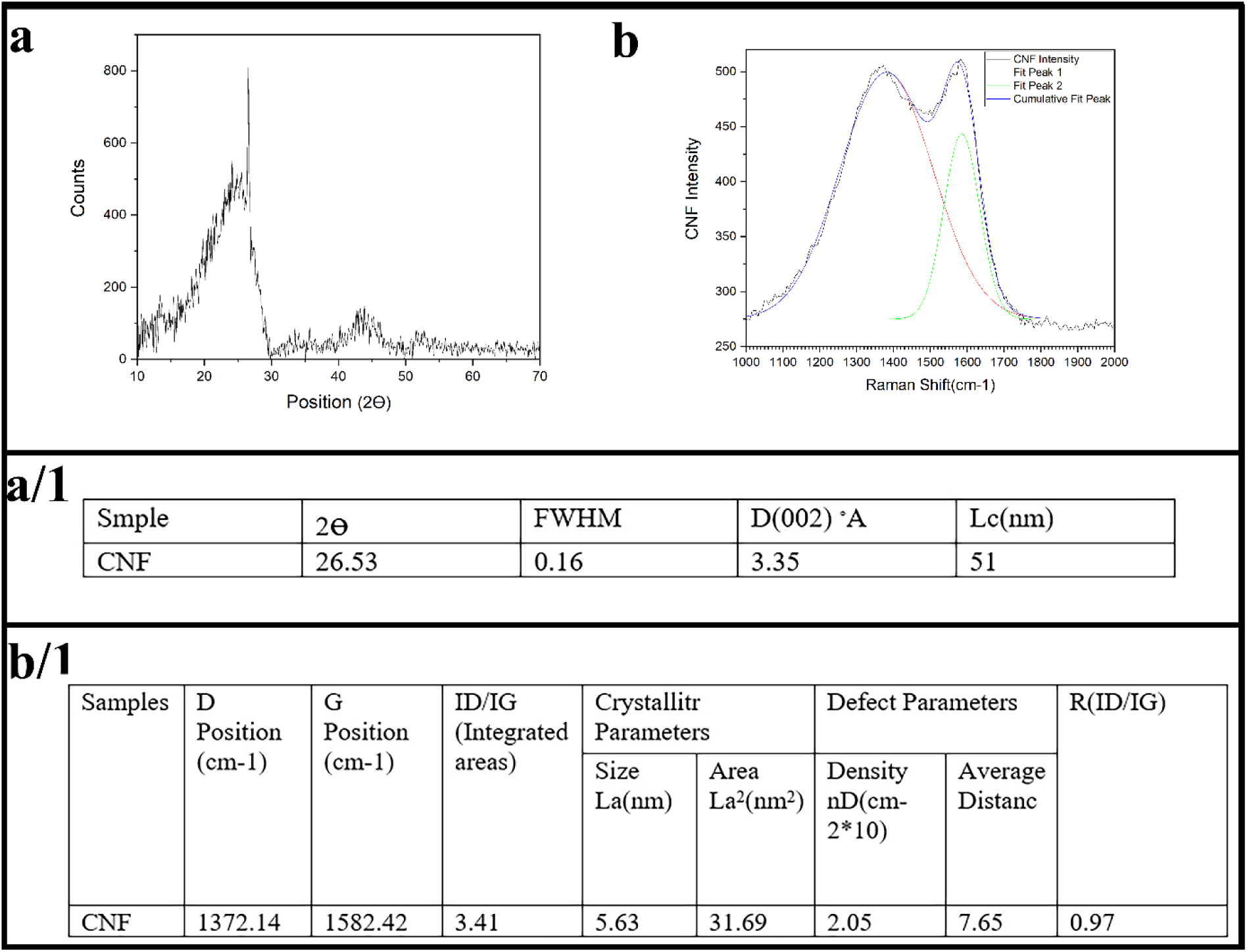
XRD and Raman results of CNFs

The hydrophobicity of the scaffold was measured with contact angle wettability investigation, which was around 110±10. This result indicates that the surface of the scaffold is highly hydrophobic.

The electrical conductivity of the scaffold was 2.87(S.cm-1) . Carbon nano fiber was prepared and characterized under the conditions mentioned in the our previous articles [52–56].

### 2.2. Isolation and characterizations of cells

Cells in passages 3 to 6 are used to perform the differentiation. hAMSC cells are spindle-shaped, elongated cells with fibroblast-like morphology. The morphology of the cells is shown in Figure 3a. In the cells cultured in the osteo and adipogenic differentiation media, the presence of colored grains in the differentiation media confirms the differentiation of cells into fat and bone by oil red and alizarin red. That approved the multipotency ability of stem cells (Fig b/1,2,3). Flow cytometry results of hAMSC cells are positive for CD90, and CD105 and negative or weakly positive for CD45, and CD34 with values in order 98%, 96%, 0.5, and 0.2 As seen in the figure 3 C. According to the results, more than 95% of the isolated cells were identified as mesenchymal stem cells.

**Figure 3:**
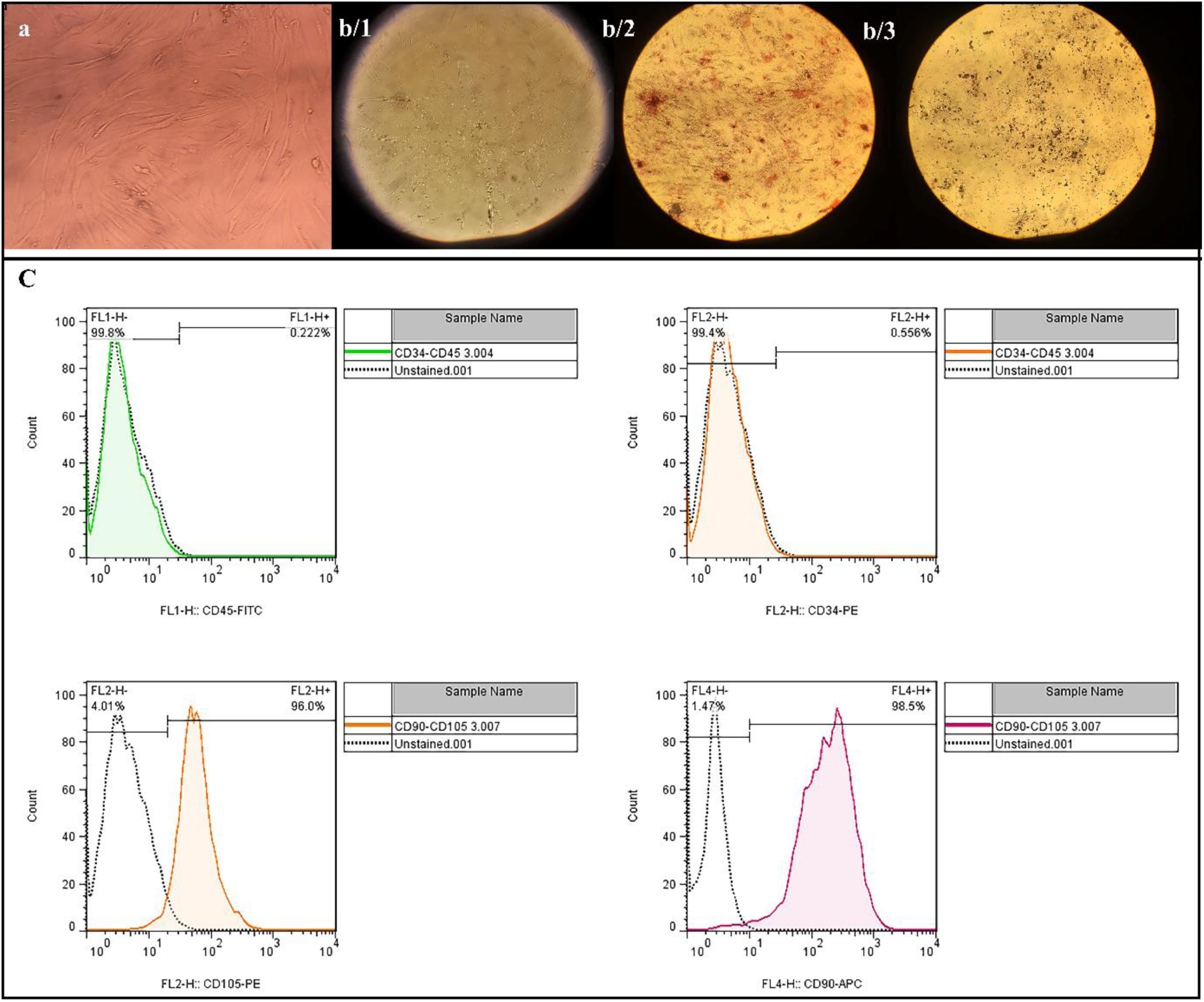
Investigation of the stem cells extracted from human adipose tissue a) observation of cell morphology b) multipotency capability of cells in fat and bone differentiation medium c) flow cytometry results for markers of mesenchymal stem cells.

### 2.3. Device design to transmit electrical current to cells

For this purpose, a structure was designed with plexiglass and 316 steel rods, which can be applied to all types of cell culture plates. The structure was autoclavable, and the rods had the necessary conductivity to transfer current. The input and output rods were connected with the help of wires so that the current entered all the cells simultaneously and equally (Figure 4). Input current intensity, frequency, and waveform are the parameters related to the electric current in each step, the optimal value was selected according to the increase in the expression of nerve cell genes and the condition of cells after electric shock.

**Figure 4:**
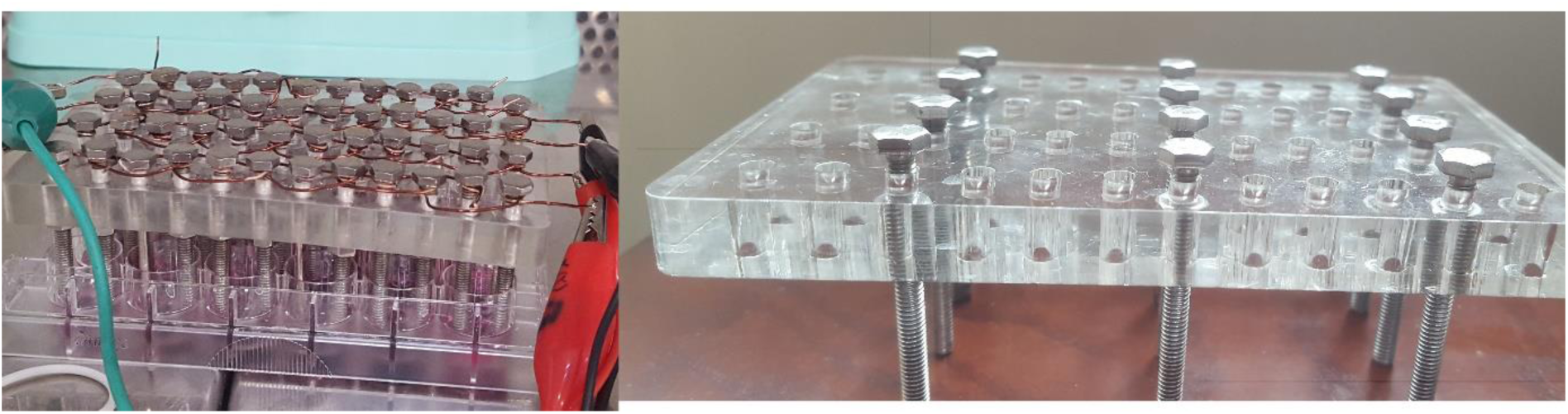
The set-up prepared for the simultaneous transfer of electric current to the cells under the same conditions.

#### 2.3.1. current intensity

Input current intensity was selected as the first parameter. Electrical stimulation of cells started from 50 uA intensity. In the other step, the current was increased at short intervals, and the frequency remained constant at 100 Hz. The cell is cultured on the scaffold, and the current is applied to the scaffolds/cells in 10 minutes for 7 days. Then the expression of nerve genes (Nestin, Map2, Tub B3, GFAP) was measured by a Real-time PCR test. higher intensities of 600, 800, 1500, and 2000 uA were applied to find the most effective current intensity, which again showed an increase in the expression of neural genes compared to the negative control (stem cells). In cells under 800 uA Compared to the 300uA, nestin expression decreased, while Map2 expression increased, which indicates the progress of the differentiation process. At 2000 µA, some attenuation was observed in the genetic content quality of the cells, so the 1500 µA current intensity was selected as the optimal current at this stage. Figure 5b shows the morphology of cells under electric current. In Figure 5, the elongated morphology of the cell confirms the change towards a neuronal cell.

**Figure 5:**
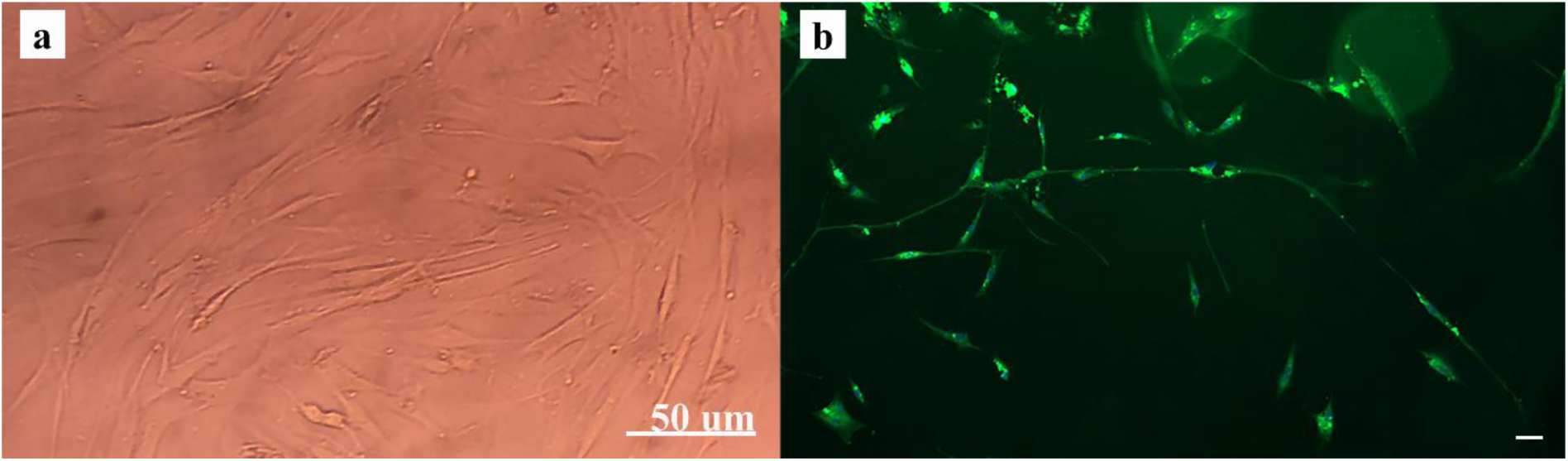
Changes in the morphology of cells during the application of electric current. a) Cell morphology before electric shock b) Cell morphology after electric shock in the form of CMOS). Figure b shows the elongation of the cells in the aftermath of an electric shock, which is a characteristic of nerve-like cells.

Figure 6 shows the Real-time PCR diagram of electrically stimulated neural genes, and increased expression of GFAP, Tub B3, and Map2 genes expression is seen in almost all groups, and there is a significant difference between the expression of genes compared to the negative control group (hAMSC). However, a decrease in nestin expression is seen during the differentiation process.

**Figure 6:**
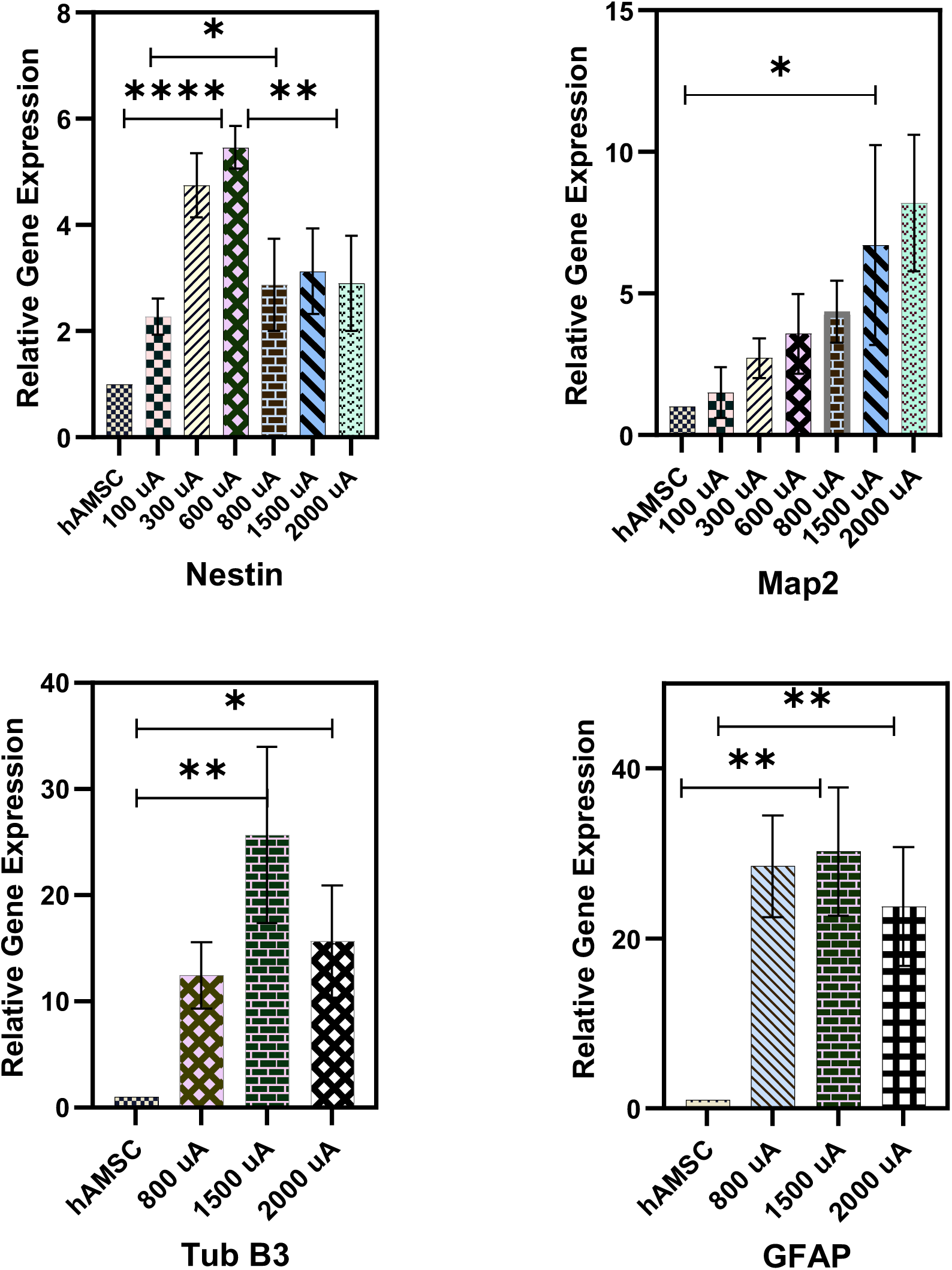
Comparison of neural gene expression in different electrical currents. Increased current intensity led to increased expression of neuronal genes

Figure 7 shows the results of nestin marker immunofluorescence in cells under electric current and its comparison with negative control (hAMSC cells) and positive control (SH-SY5Y cells).

**Figure 7:**
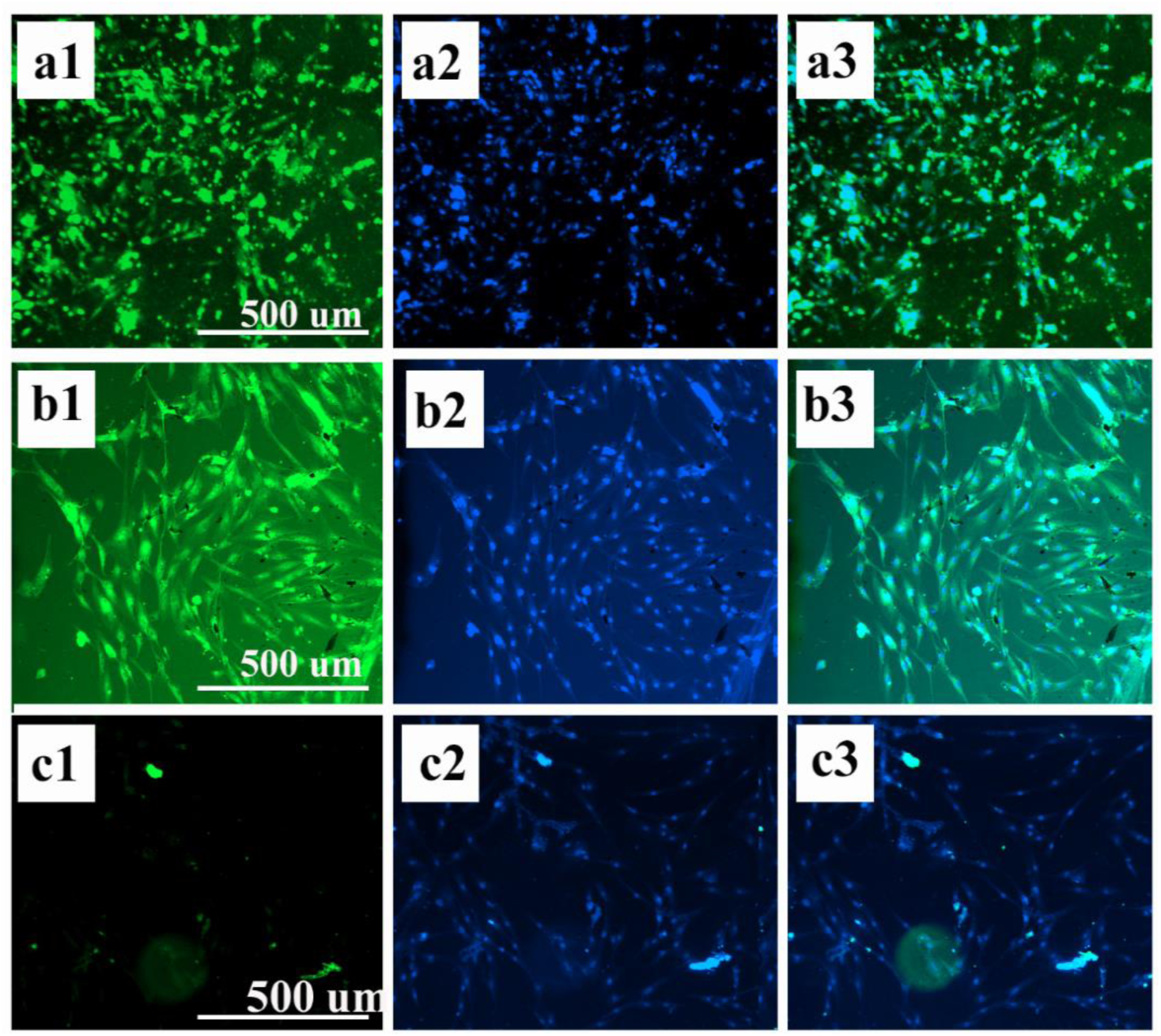
Fluorescence staining for nestin marker in a, b, c) positive control (SH-SY5Y cells) d, e, f) cells under electric current for seven days and 10 minutes g, h, i) negative control, hASMC stem cells

#### 2.3.2. Optimal frequency selection

Four frequencies of 100, 500, 1000, and 2000 Hz were investigated at the current intensity of 1500 uA in this study. The other parameters of current intensity, waveform, shock time, and the number of shock days of stimulation remain constant. At 2000 Hz, cell deaths were observed from day 4, so the 2000 Hz group was excluded. Examination of Real-time PCR results in Figure 8 by comparison between the three groups showed that the frequency of 500 Hz showed a better result than the other groups.

**Figure 8:**
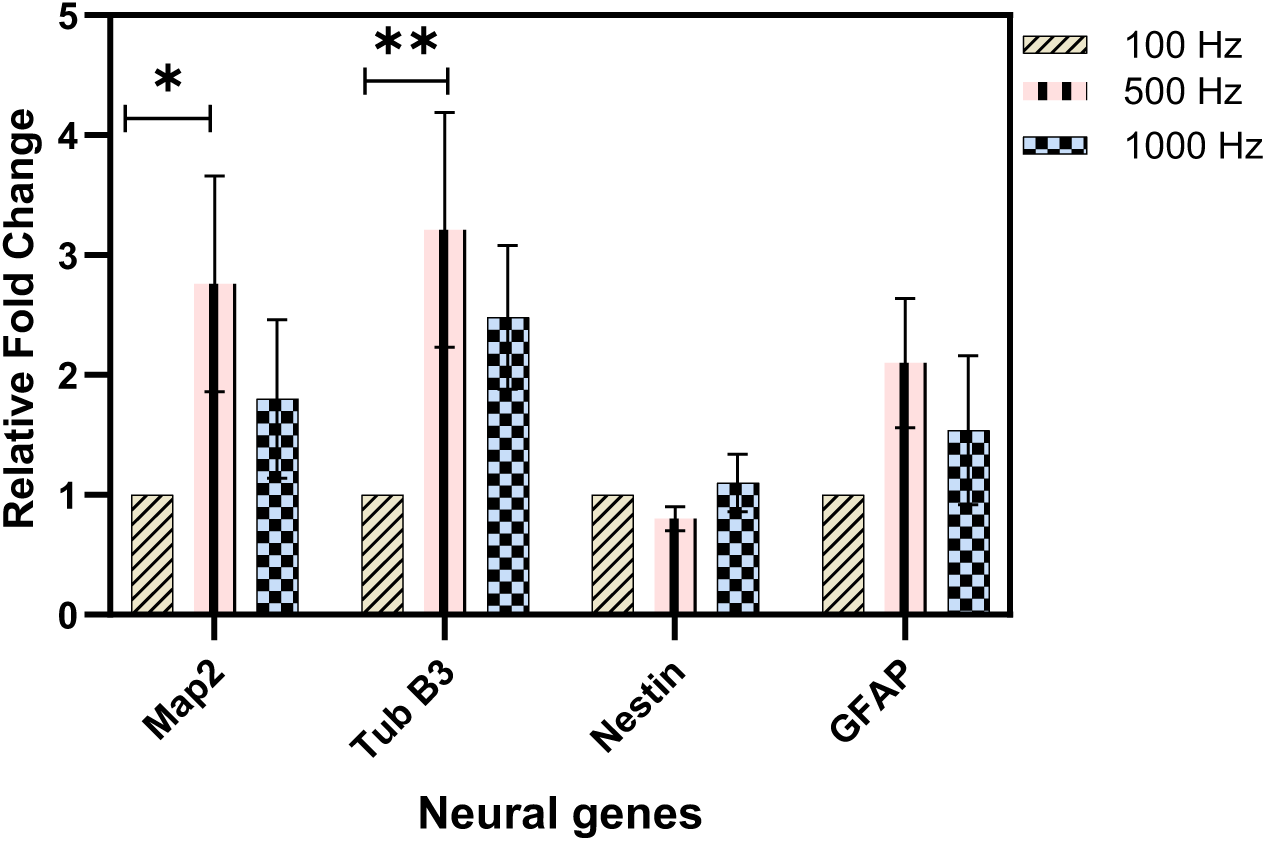
Comparison of gene expression at different frequencies. A significant increase in Map2 and tubulin expression was seen at 500 Hz.

#### 2.3.3. Select the appropriate waveform

In this section, three forms of CMOS, sinusoidal and square were used. In comparison between these groups, different results were seen in the morphology of the cells. In the sinusoidal and square waves, the morphology of the cells was different from that of the CMOS form, as shown in Figure 9. On day 8, most of the cells were abruptly destroyed but saw a lot of morphological changes in the cells on the final before cell death, so the morphological changes were greater than in the CMOS model. It seems to have more differential power in sinusoidal and square form, but due to the loss of cells, it was not possible to continue the study for more days and longer periods of shock. The comparison between square and sinusoidal shows that the square form shows more expression of neuronal gene markers. Figure 10 shows real-time PCR results between different groups of waveforms.

**Figure 9:**
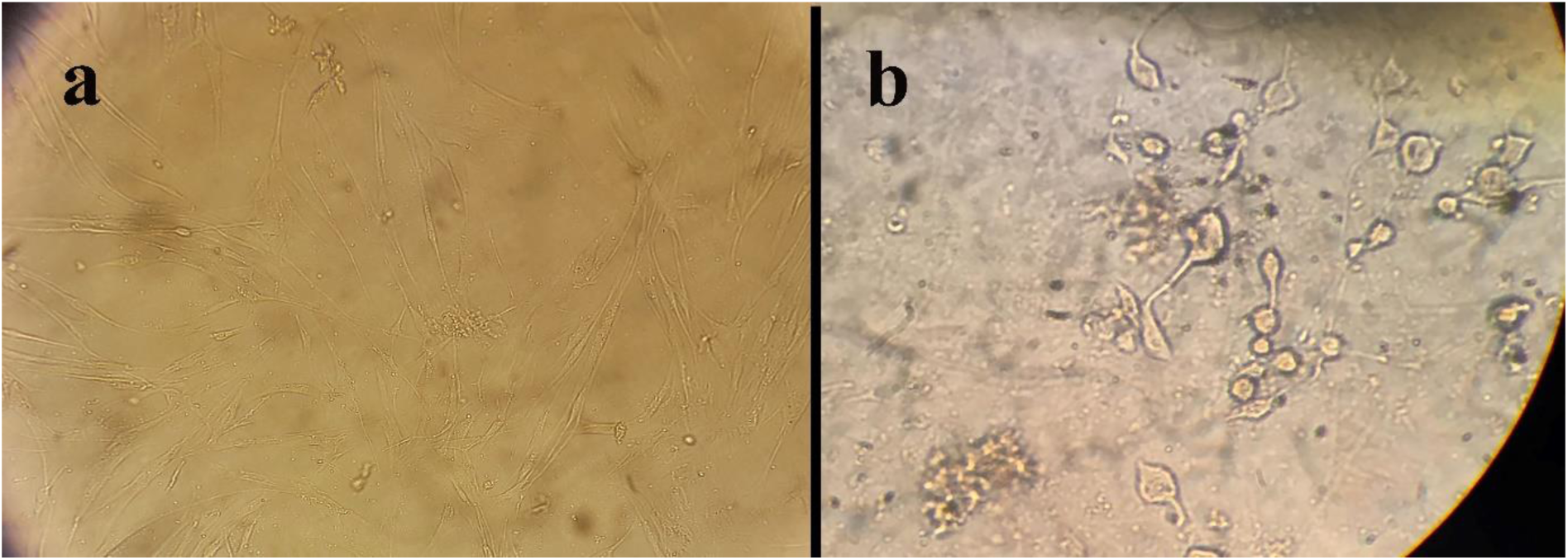
a) hAMSC cells without electric shock treatment (magnification ×20), and b) cell morphology after 6 days of electric shock with square waves (magnification ×40)

**Figure 10:**
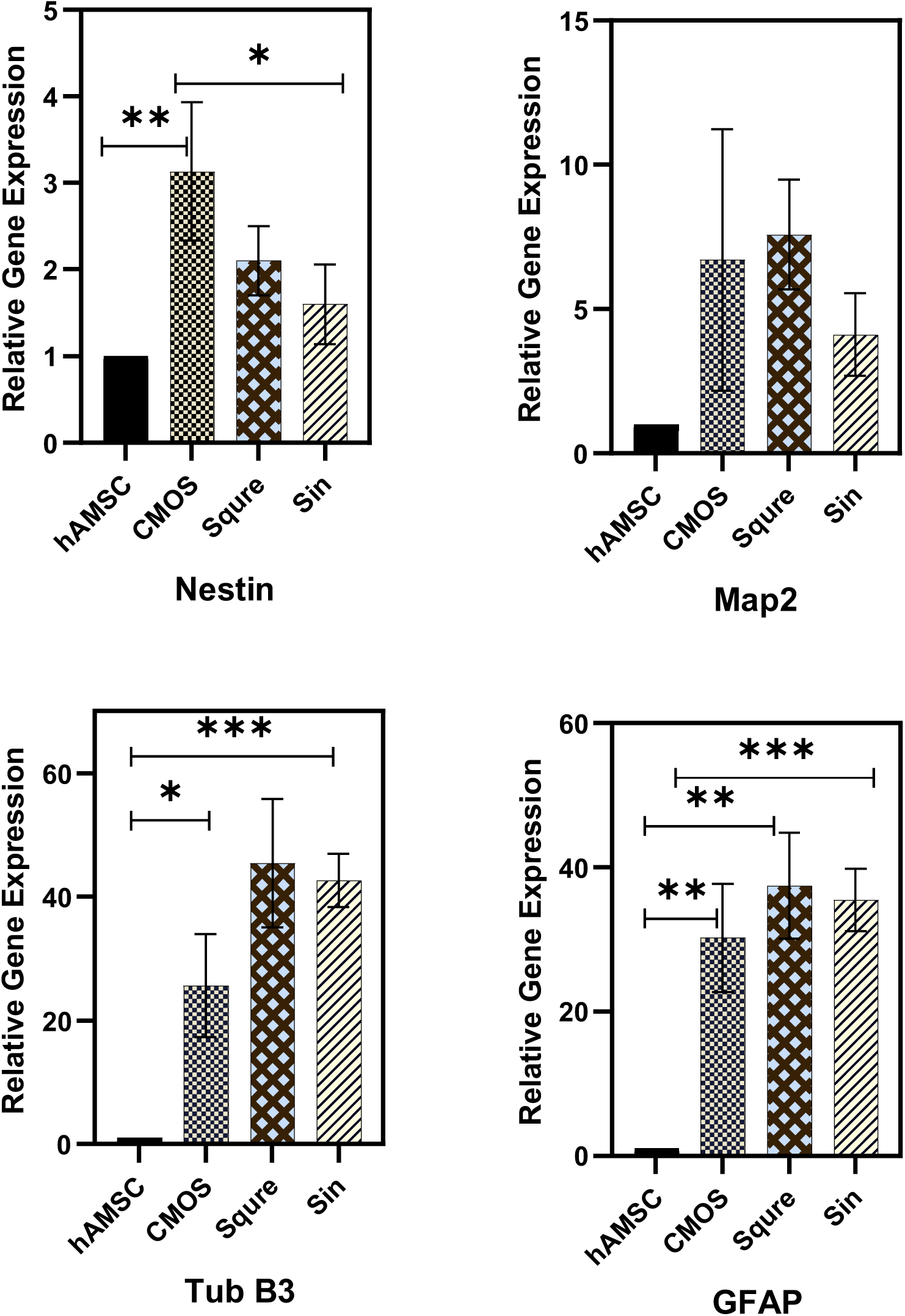
Comparison of gene expression in different waveforms. In sinusoidal and square forms, there is an increase in the expression of nerve genes.

#### 2.3.4. Select the daily duration of the shock

The optimal electrical flow parameters were selected in the previous steps. 1500 µA current and 500 Hz frequency with CMOS waveform were selected, and the cells were exposed to currents in the daily times of 10, 30, and 60 minutes. Real-time PCR results in Figure 11 show that by enhancing the shock time, more expression of neural genes is seen in the cells. The difference between 30 and 60 minutes is not significant, but there is a significant difference with 10 minutes, and at 60 minutes we see a decrease in GFAP, which can be said that by increasing the shock time, the expression of neuronal to glial genes improves.

**Figure 11:**
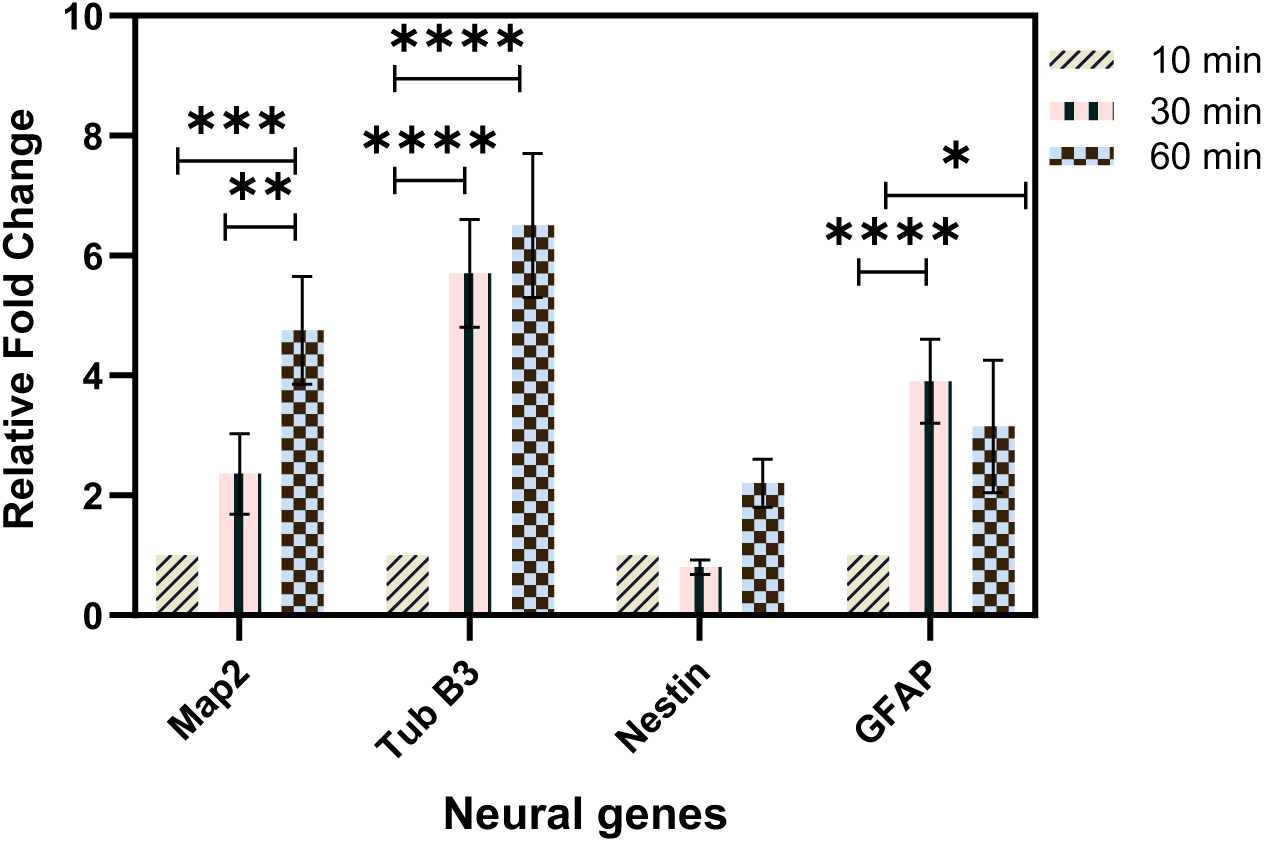
Comparison of gene expression due to changes in daily shock time. With increasing shock time, an increase in neuronal markers is observed.

#### 2.3.5. Optimal stimulation duration

The duration of electrical stimulation was studied as another factor. Differentiation was performed with electric shock for 7 days, 14 days, and 21 days. Optimal factors include 1500 uA, 500 Hz, and CMOS waveform were selected. As shown in Figure 12. It was observed that with increasing the duration of the shock period, more maturation neuronal markers were seen compared to the GFAP. increase in Map2 indicates that the cells progress to maturity as the length of the period increases .Figure 13 shows the immunofluorescence staining of cells at different periods for the nestin marker.

**Figure 12:**
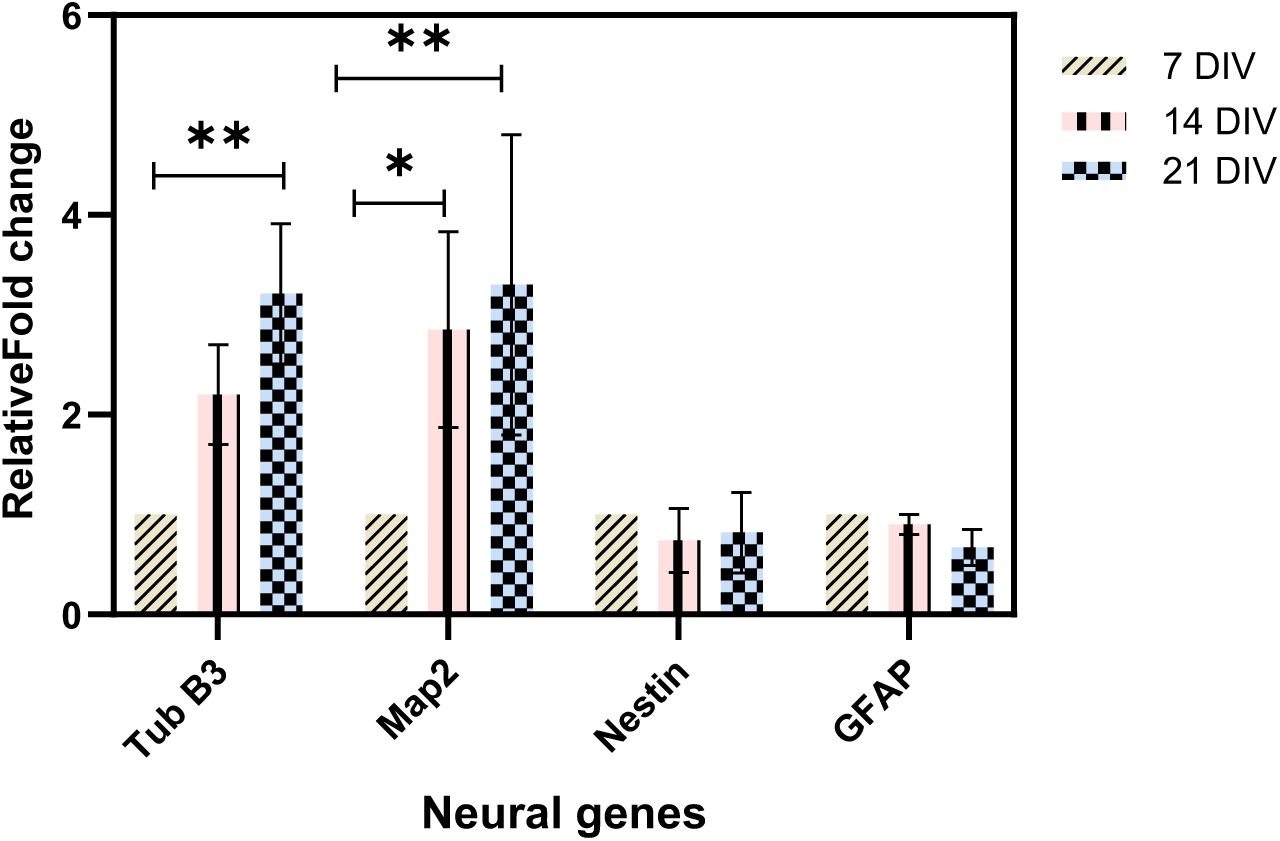
Comparison of the expression of neuronal genes during different periods of shock, which decreased with increasing length of shock period, decreased GFAP marker, and increased neuronal markers.

**Figure 13:**
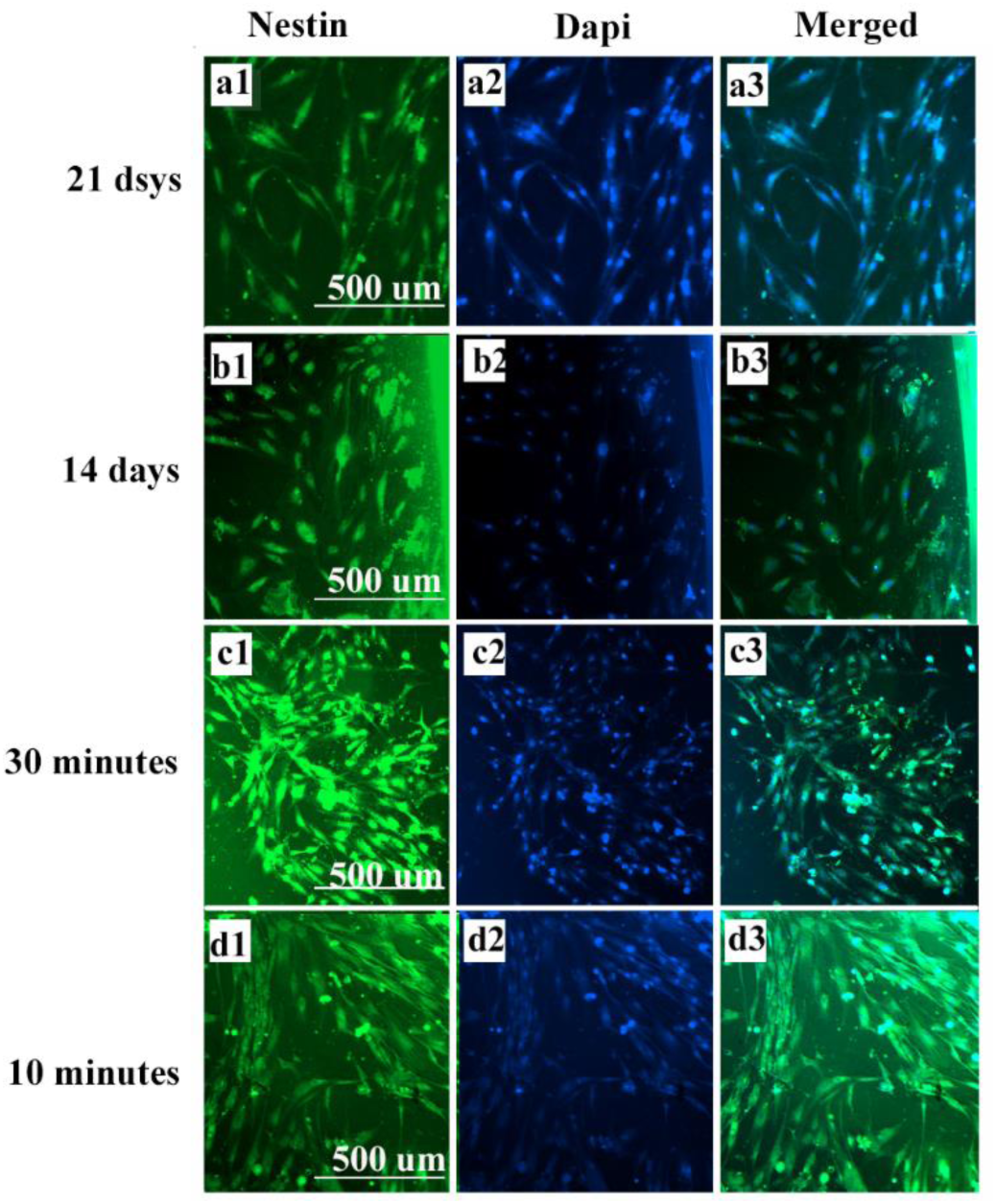
Nestin immunofluorescence in periods and times of shock nestin marker was seen in all groups under electrical stimulation.

Overall studies show that the current intensity of 1500 uA, with a frequency of 500 Hz, with a square biphasic waveform shows the expression of more neural genes. Time studies show that the longer the daily shock time, the higher the gene expression, and this test is performed for up to 30 minutes. Another study shows that increasing the duration of the shock period to 14 days and 21 days shows a significant decrease in the expression of GFAP as a glial factor. Therefore, by adjusting the current parameters, it is possible to differentiate hAMSC into neurons like cells in DMEM High Glucose + 10% FBS medium. In Figure 14, there is an immunofluorescence image of the cells in selected conditions of each parameter and shows the expression of Map2 and Tub B markers.

**Figure 14:**
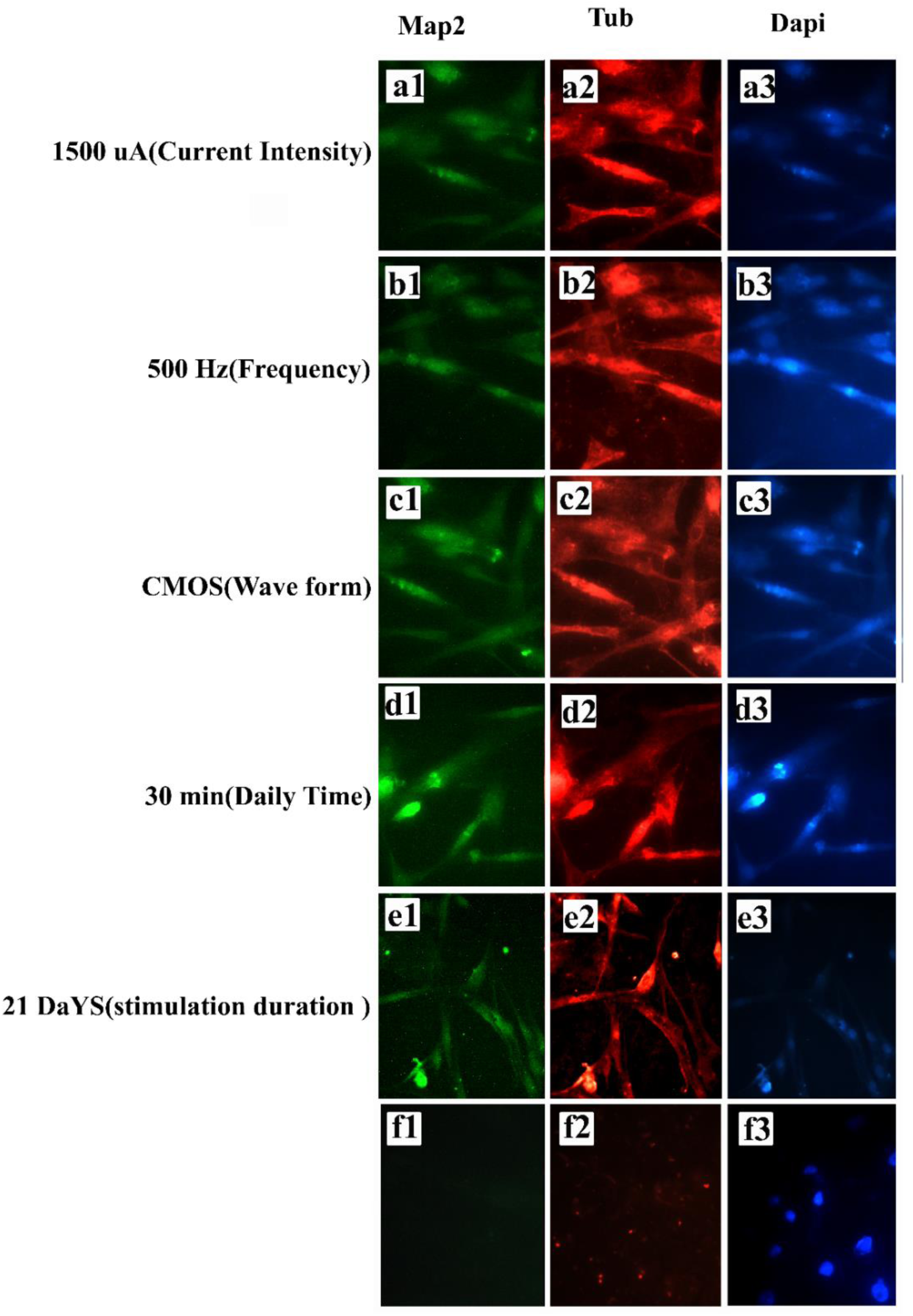
Immunofluorescence images of Map2, Tub B3 cell markers in optimal conditions of each electric current parameter.

### 2.4. Investigating the effect of scaffold presence on gene expression

Real-time PCR results for electrical stimulation in the presence and absence of CNF scaffolding are shown in Figure 15. The results show an increase in the expression of neural markers in the presence of carbon nanofibers. Therefore, CNF has been effective in increasing the expression of nerve genes .It seems that the electrical conductivity of the scaffold and the nanofibrous shape of the scaffold affected the neuronal differentiation of the cells.

**Figure 15:**
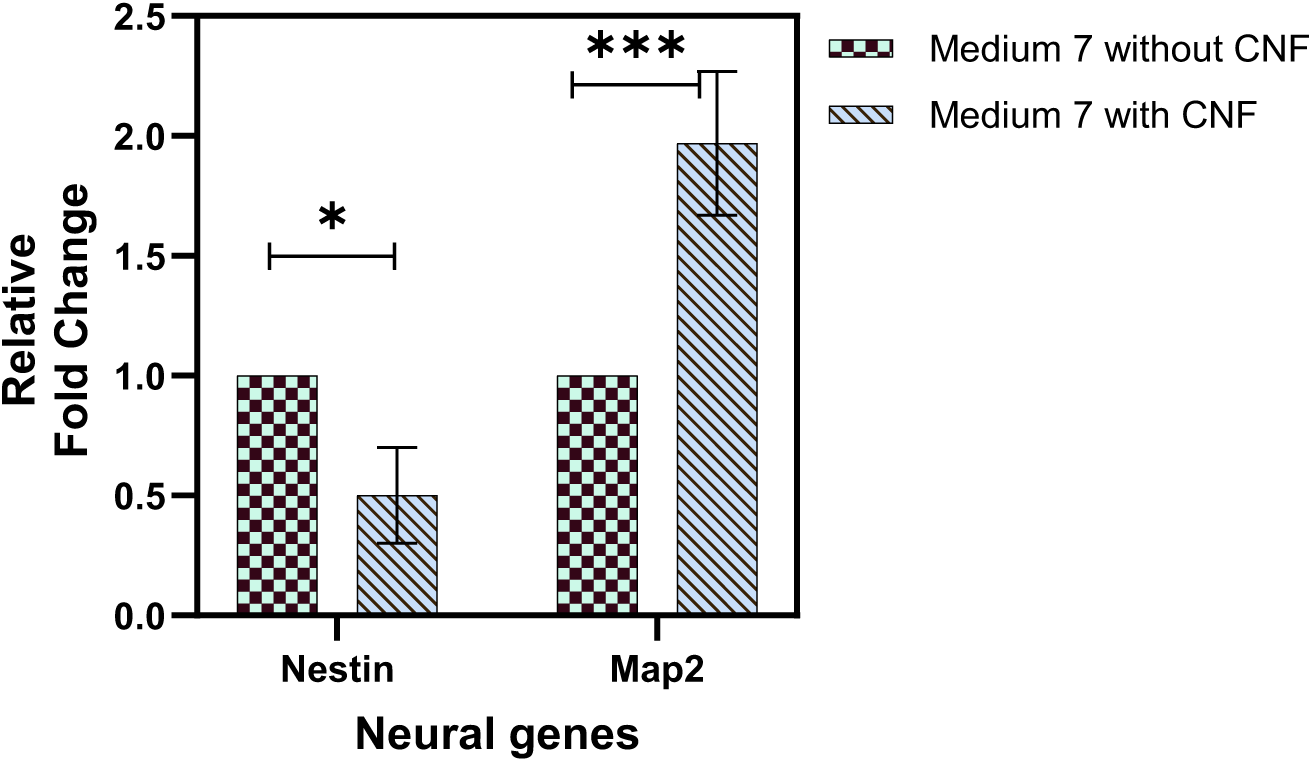
Comparison of gene expression in the presence and absence of CNFs. The results show that the presence of CNFs is effective in differentiation under electrical stimulation.

### 2.5. hAMSC differentiation in nerve differentiation medium

Cells were cultured on CNF scaffolds and pre-differentiation and differentiation media were added to the cells according to the stated sequence. Seven days later, RNA cells were extracted and the results of real-time PCR were evaluated. Figure 16 compares the expression of Map2 and nestin genes on day seven. In this part of the study, the expression is compared to the stem cells (Fig 16a). Figure 17b the differentiation medium in the presence and absence of CNFs with the chemical medium. It shows that in the differentiation medium, carbon nanofiber increases the intensity of differentiation, conductive scaffolds can be effective for the differentiation of nerve cells in the absence of current, and in addition, the shape of the nanofiber can also be effective. Figure 16c the comparison between the differentiation with electrical stimulation and chemical differentiation during 7 days. It shows that differentiation with differentiation medium shows more expression of neural markers than differentiation with electrical stimulation. Cells were immobilized and immunofluorescence of nestin, Map2, and Tub B3 was evaluated. Figure 17 Cells were stained with nestin and DAPI.

**Figure 16:**
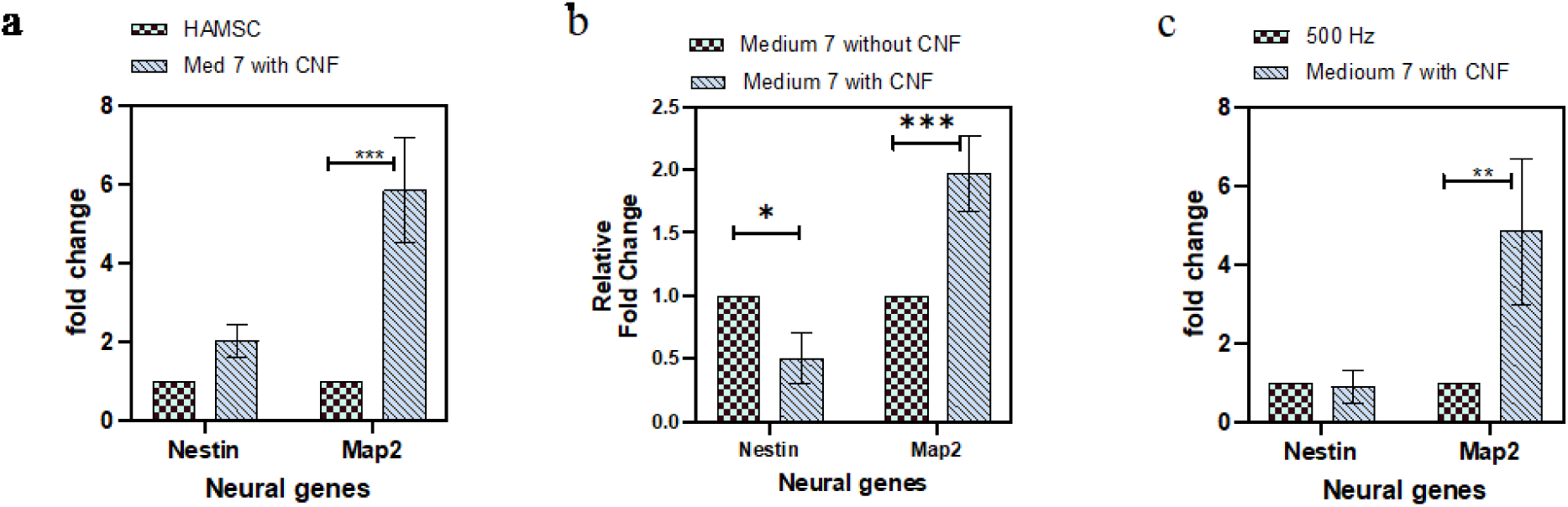
Results of neuronal gene expression in differentiation and electrical stimulation groups. Carbon nanofibers are effective in increasing neuronal differentiation chemically and there is a significant difference.

**Figure 17:**
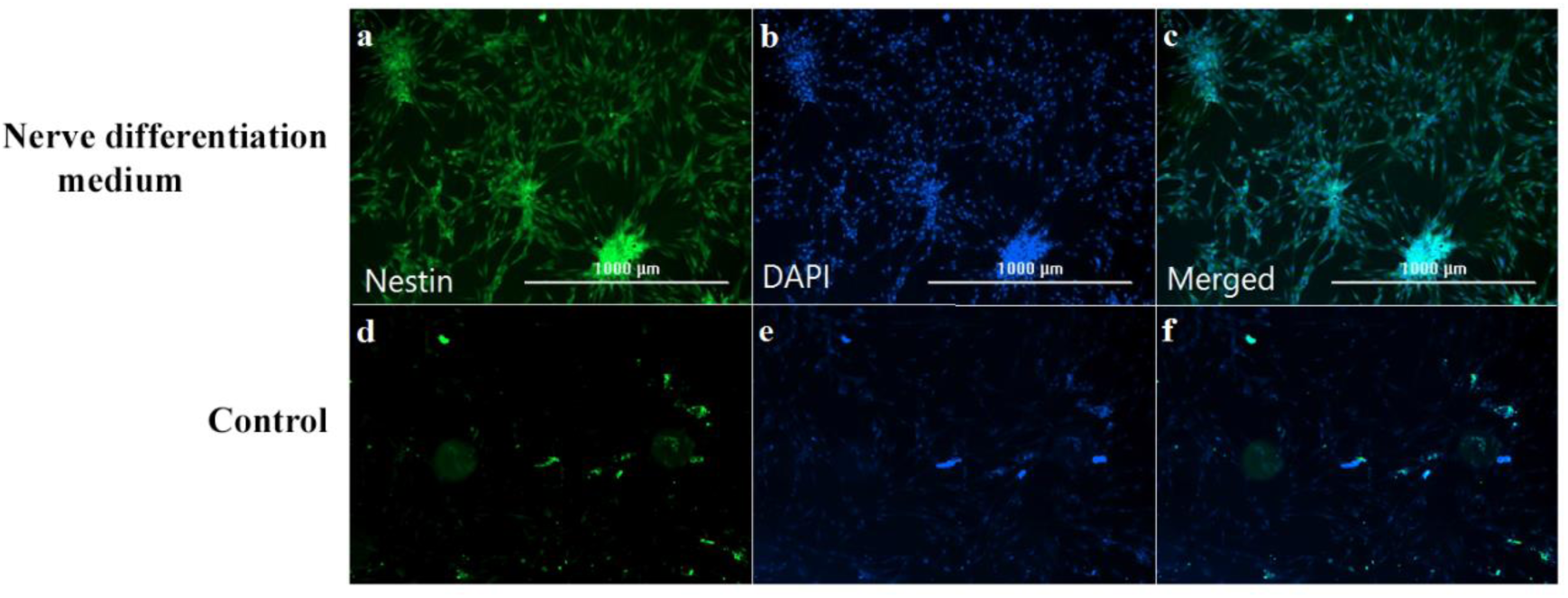
Immunofluorescent staining with Nestin antibody and DAPI differentiated cells with neural differentiation medium

### 2.6. Investigation of change in stem cell surface markers

When stem cells enter the differentiation phase, specific surface markers of these cells, such as CD105 and CD 90, show a decrease in expression, and by flow cytometric testing, a significant decrease in the expression of positive and specific markers of Mesenchymal stem cells CD90 and CD105 after electrical stimulation was seen. The CD90 marker decreases from 98% to 85% of expression, and the CD105 marker decreases from 96% to 73% of expression. The results are shown in Figures 18.

**Figure 18:**
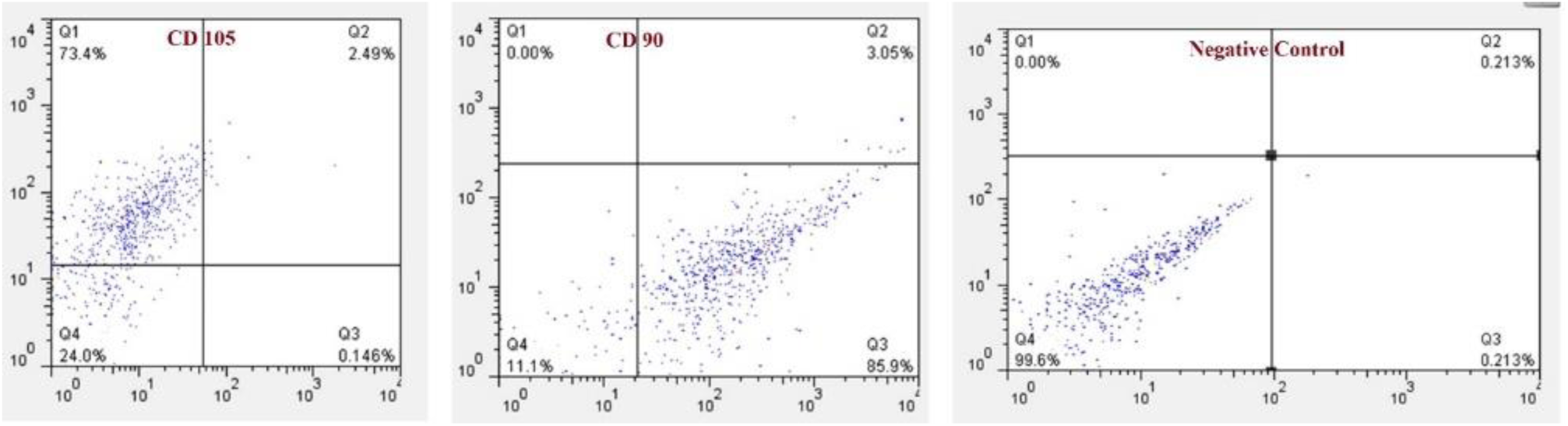
Flow cytometric diagram of cells after electric shock and decreased expression of stem cell markers. Decreased expression of specific stem cell markers confirms differentiation.

### 2.7. Study of cell migration

In this study, cells were cultured in plates around the wells. the CNF scaffold was cut to equal size and placed in the center of the well. Two wells were under electrical stimulation, and two were without electrical stimulation. Two wells without CNF and electrical stimulation as control. After 10 days of applying the current with the adjusted parameters, it was observed that the cells moved toward the scaffold, but no difference was seen in the model with and without electric current (1500 uA, 500 Hz, CMOS, 10 minutes per day for 7 days), as shown in Figure 19. According to the articles, electric current is involved in cell migration. It seems that the electric current needs its parameters to migrate and the current intensity must be adjusted. Many papers have suggested that direct current has a greater effect on cell migration than alternating current.

**Figure 19:**
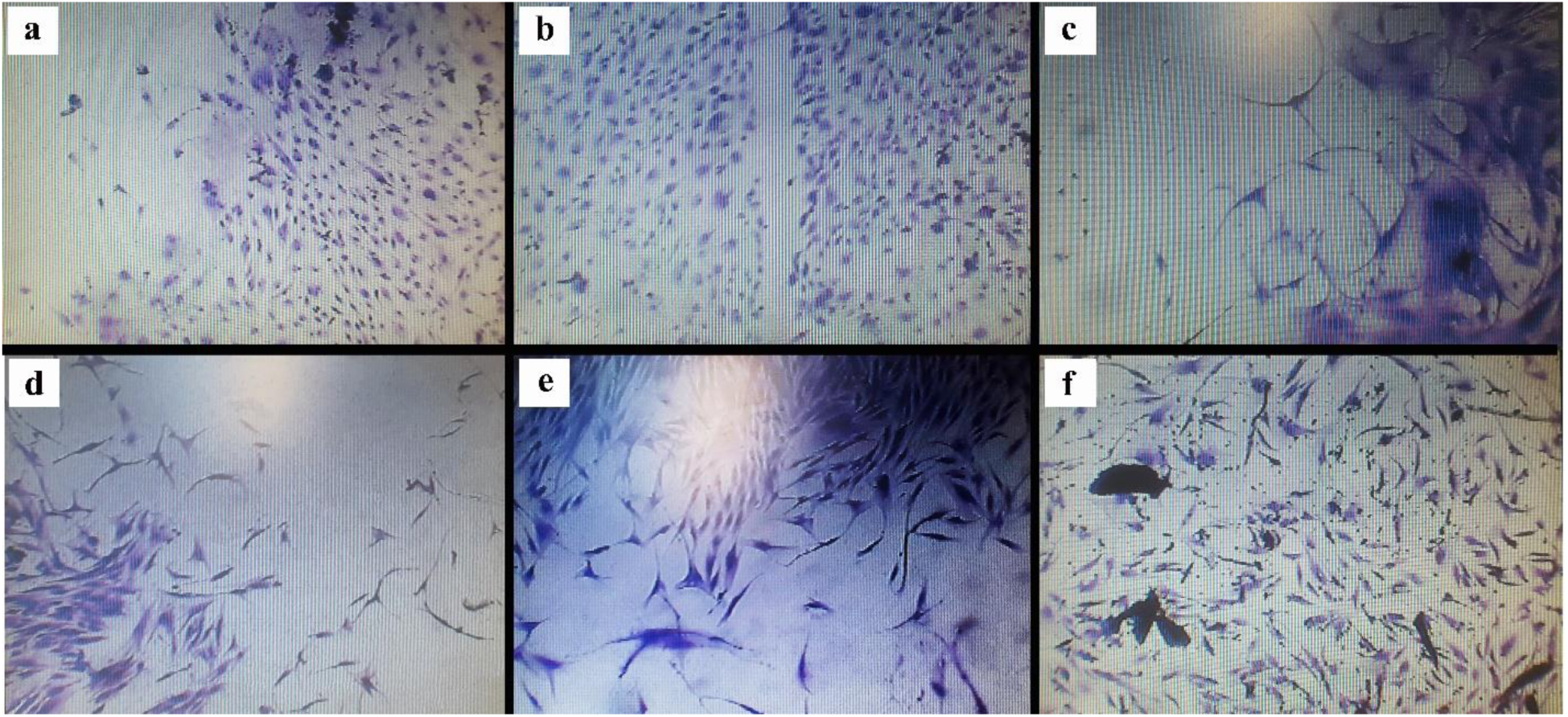
Migration of stem cells in the presence of scaffolding and electric current a, b) Control without scaffolding and with electric current (Magnification 10X) c) Cells in the presence of scaffolding and electric current 4 d) Cells on day 6 in Presence of current and scaffolding e) on day 8 in the presence of current and scaffolding f) on day 10 in the presence of scaffolding. Carbon nanofibers cause cells to migrate to the scaffold. c, d, e, f: Magnification: 20X))

**Figure 20:**
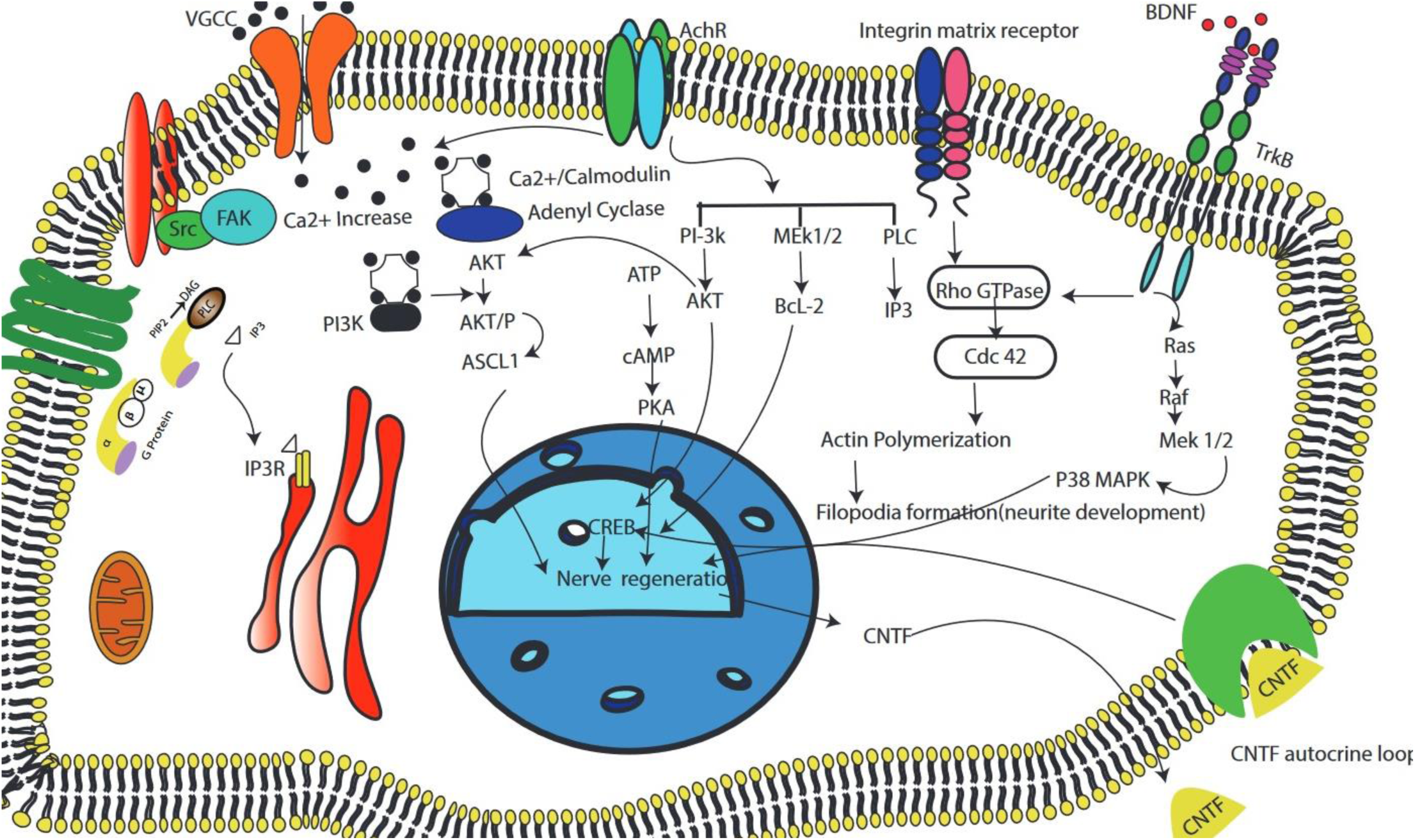
Schematic view of cellular mechanisms effective in differentiation activated by electrical stimulation.

## 3. Discussion

Neurodegeneration and traumatic injuries to the brain or spinal cord are associated with various complications, and the prevalence of these diseases increases every year [57]. One of the treatment methods is based on cell therapy. Adipose tissue-derived cells (ADSCs) are a good choice because they are abundant in the body, and the ethical challenge is minimal. Also, extracting these cells does not cause trauma in the body, and ADSCs cells can differentiated into neurons [58]. In this research, the differentiation of mesenchymal stem cells derived from human adipose tissue into nerve cells was investigated under electrical stimulation and without environment and differentiation factors.

Different studies have demonstrated that scaffolds with nanofiber morphology and electrical conductivity are favorable for the differentiation and development of nerve cells [59]. These reports exhibit that nanoscale topography synergistically improves the expression of neural markers and electrophysiological characteristics and functional maturation of neurons differentiated from human NSCs [60, 61]. Nano or micro-topography factors affect different cell mechanisms from changing cell morphology, to cell skeleton, and cell biomechanical changes, which leads to the start of signaling cascades [62, 63]

In recent studies, due to the problems of differentiation with chemical environments, more attention is paid to the approach of using physical stimuli. One of these stimuli is the use of electronic stimulation, and different parameters of the electric current and the type of cell used can influence the results. In an alternating electric field, cell differentiation can be the result of the frequency of the field. In this study, the effect of different frequency values on mouse NSC cells is studied [64].

Electrical stimulation can be monophasic or biphasic. Biphasic may be more useful because it prevents charge accumulation produces fewer electrolysis products at the electrodes, minimizes heat generation, and increases signal transmission. It can be applied over longer time intervals and at higher voltages. For this reason, biphasic stimulation is often used for clinical applications of nerve tissue [64, 65]. In the comparison between sinusoidal and square waveforms, the square waveform showed more gene expression. This result is probably because of the gradual changes in the current in the sinusoidal waveform compared to the square waveform, where the current suddenly changes from positive to negative.

Chang et al. reports demonstrated that biphasic electric current causes neuronal proliferation and differentiation by reducing glial factor from NPCs in mouse embryos [65, 66]. Expression of GFAP is related to an increase in the concentration of calcium and STAT3 [67].

In a series of studies, electrical stimulation on NPCs of mouse embryos on an alginate hydrogel scaffold shows an increase in astrocyte differentiation compared to neuronal differentiation at a certain frequency. The results show that alternating electric fields can manipulate the differentiation to generate immature cells, which is valuable for future clinical applications [68, 69]. Lim et al. showed that electrical stimulation changes the ratio of neurons and astrocytes, but no oligodendrocytes were observed. This study also showed that high-frequency electrical stimulation delays the differentiation into mature astrocytes. The response to electrical stimulation depends on the cell type, developmental stage, and diverse species [69].

It seems that the mechanism of differentiation activated by topography and electrical stimulation is by activation of the b-catenin/Wnt signaling pathway [70–72]. Mao et al. stated that in addition to chemical factors, topography could affect the Wnt cell signaling [73]. On the other hand, it is mentioned in the articles that the increase of reactive oxygen groups activates the b-catenin/Wnt pathway [74]. On the other hand, one of the important factors that activate this signaling pathway is the increase in the amount of calcium released from the cells [28], and in this study, the effect of the current on the cellular pathway is also stated [75], of course, confirmation of this issue requires the investigation of molecular pathways with special tests. The effect of electrical stimulation on the regulation of neural differentiation depends on the cell type. The sensitivity of mesenchymal cells under an electric field is more than NSC [60, 67].

The cell type and its development stage effect on the differentiation process was predicted in the previous research. It was reported that the characteristics of the cells were different depending on which part of the separated fat was applied [28]. In most of the studies in which neural differentiation has been done with electric current, NSC cells have been applied, and they have been used in the neurobasal environment or together with growth factors. These growth factors themselves are effective in the differentiation of nerve cells. In this study, mesenchymal stem cells and DMEM High Glucose + 10% FBS basic medium were applied, and no growth or differentiation factors were applied. A very long duration of stimulation or high field strength causes negative effects such as shortening the length of neurons or indeterminate morphology. Figure 21 shows the mechanisms identified in the differentiation of stem cell to nerve cell using electrical stimulation.

Electrical stimulation with longer shock periods and shorter daily shock time causes neural differentiation and maturation of cells. Unlike DC, the AC does not affect the migration and lineage of NSC cells. The reason for this can be the bidirectional electric field resulting from the alternating field. According to the results, it seems that with the passage of time and the continuation of electrical stimulation, the cells show a greater tendency to become neurons than glial cells.

## 5. Conclusion

Our results indicate that carbon nanofiber is a suitable microstructure in terms of electrical and topographic conductivity for nerve studies. Stem cells have a high affinity for carbon nanofiber structure, and the migration of cells to the scaffold is observed. Further functionalization of carbon nanofibers leads to improved cellular adherence, proliferation, and differentiation [52]. Examination of the nerve gene expression by electrical stimulation demonstrates that this method can direct the neuronal differentiation of cells to an appropriate level. Carbon-based materials have excellent electrical properties and are easily biofunctionalized. Therefore, we used carbon-based nanofiber scaffolds to make them electrically conductive. From 100 to 2000 µA, we had an increase in the expression of neural markers. At low current, the expression of nestin and Map2 increased, and with further increasing current, it was observed that the expression of nestin decreased over time, and the expression of Map2 increased. This rhythm of gene expression is seen during the differentiation of nerve cells since nestin is considered a progenitor marker and Map2 is seen in mature nerve cells. The reduction of nestin expression occurs in the progress of the differentiation process. After determining the optimal intensity of electric current, we investigated the effect of different frequencies.

The waveform describes the shape of one cycle of the voltage or current. In the next step, the sinusoidal and square waveforms were investigated. The change in the morphology of the cells in this method was very striking. However, the glial marker expression was more than the CMOS samples, and on the other hand, the stability of the cells was much less. So, on the 7th day, the cells suddenly died. Perhaps the increase in glial markers was due to the faster destruction of neuronal cells in the absence of specific environments containing growth factors.

In another attempt, study times were changed daily and periodically. An increase in the expression of neural genes was seen with an increase in the daily shock time. In another model, the length of the stimulation period was increased to 14 and 21 days. It was seen that with the increase in the duration of the period, the expression of mature neural markers increases, and a decrease in progenitor markers is seen, which shows that by increasing the period of electrical stimulation, the percentage of neuronal to glial cells can be increased. The summary of the results of different electric current parameters is stated in Table 2.

**Table 2:** The conditions of applying electrical stimulation in groups and the optimal results of different parameters of electrical current.

| Variable Parameter | The Values of variable Parameter | Constant Parameter | Best Results |
| --- | --- | --- | --- |
| Current Intensity | 100, 300, 600, 800, 1500, 2000 uA | Frequency: 100 Hz<br>Wave form: CMOS<br>Daily shock duration: 10 min<br>Culture Medium: Dmem Hg+ 10% FBS<br>Duration of stimulation: 7 days | 1500 uA |
| Frequency | 100, 500 , 1000 | Current Intensity: 1500 uA<br>Wave form: CMOS<br>Daily shock duration: 10 min<br>Culture Medium: Dmem Hg+ 10% FBS<br>Duration of stimulation: 7 days | 500 Hz |
| Wave forms | CMOS, Sinusoidal, Square | Current Intensity: 1500 uA<br>Frequency: 500 Hz<br>Daily shock duration: 10 min<br>Culture Medium: Dmem Hg+ 10% FBS<br>Duration of stimulation: 7 days | Sinusoidal, |
| Daily shock duration | 10 , 30 , 60 min | Current Intensity: 1500 uA<br>Frequency: 500 Hz<br>Wave form: CMOS<br>Culture Medium: Dmem Hg+ 10% FBS<br>Duration of stimulation: 7 days | 30 min |
| Duration of stimulation: | 7, 14 , 21 days | Current Intensity: 1500 uA<br>Frequency: 500 Hz<br>Wave form: CMOS<br>Daily shock duration: 10 min<br>Culture Medium: Dmem Hg+ 10% FBS | 21 days |

In another model, the effect of current in the presence and absence of scaffolding was investigated with electric current, and the presence of scaffolding showed an increase in the expression of neural markers compared to its absence. Of course, in the group without scaffolds, an increase in the expression of neural markers was seen compared to the negative control group, and it showed that even without scaffolds, neural differentiation with electrical stimulation. It happens because of the conductivity of the culture medium.

Real-time PCR results indicated that until the 8th day of the test, significant expression of GFAP, equal to tubulin was observed, considering that both tubulin and GFAP genes are among the starting genes in neural differentiation. However, in the increase in the number of days of electric shock, a decrease in the expression of GFAP and an increase in MAP2 and tubulin were seen.

Therefore, this method can be replaced by differentiation techniques based on chemicals and growth factors due to their limitation such as the high cost, low accessibility, and low stability. The toxicity of substances in chemical differentiation, including DMSO and 2Me, has limited the use of these substances. One of the advantages of the current method based on electrical stimulation is that it can be disconnected or connected at any time. Electrical stimulation can be applied to any part of the scaffold.

## Declarations

### Ethics approval and consent to participate

This study received ethics approval from Research Ethics Committes of School of Medicine-Tehran University of Medical Sciences, Approval ID: IR.TUMS.MEDICINE.REC.1400.1531

### Consent for publication

Not applicable

### Availability of data and Materials

All data are presented in the article

### Competing interests

The authors confirm that there are no financial or commercial relationships that lead to Conflict of interest

### Funding

This work was supported by Tehran University of Medical Sciences, grant no. 98-01-87-41015.

### Author’s Contributions

The authors confirm contribution to the paper as follows: study conception and design: Houra Nekounam, H.Golmohammady; data collection: H.Nekounam,H.Golmohammady; analysis and interpretation of results: H.Nekounam, MA.Shokrgozar; draft manuscript preparation: H.Nekounam, SM.Amini. All authors reviewed the results and approved the final version of the manuscript.

## Acknowledgements

Not Applicable

## Notes

### Competing Interest Statement

The authors have declared no competing interest.

